# Human CD1c-autoreactive T-cells recognise *Mycobacterium tuberculosis*–infected antigen-presenting cells and display cytotoxic effector programmes

**DOI:** 10.64898/2026.01.16.700025

**Authors:** Matthew Milton, Sahar H Farag, Diana Garay-Baquero, Jennie Gullick, Kinga Niedobecka, Daniel Burns, Rita Szoke-Kovacs, Patrick Trimby-Smith, Alex Look, Richard Stopforth, Marco Lepore, David K Cole, Laura Denney, Andrew White, Sally Sharpe, Alasdair Leslie, Andres Vallejo, Liku Tezera, Paul Elkington, Salah Mansour

**Affiliations:** School of Clinical and Experimental Sciences, NIHR Southampton Biomedical Research Centre, Faculty of Medicine, University of Southampton, Southampton SO16 6YD, United Kingdom; Research and Evaluation, UK Health Security Agency, Porton Down, Salisbury SP4 0JG, United Kingdom; Immunocore Limited, Abingdon, Oxon OX14 4RY, United Kingdom; Africa Health Research Institute, Durban, South Africa; School of Laboratory Medicine and Medical Sciences, University of KwaZulu-Natal, Durban, South Africa; Division of Infection and Immunity, University College London, London, United Kingdom; Institute for Life Sciences, University of Southampton, Southampton, United Kingdom

**Author notes:** **Address correspondence to**: Salah Mansour, Faculty of Medicine, University of Southampton, Clinical and Experimental Sciences, MP 813, LE 77. Southampton General Hospital, Southampton SO16 6YD, UK. These authors contributed equally to this work (Matthew Milton, Sahar H Farag).

## Abstract

Tuberculosis (TB), caused by *Mycobacterium tuberculosis* (Mtb), remains the leading cause of death from infection globally, yet the contribution of non-classical T-cell pathways to human immunity remains poorly defined. CD1c-autoreactive T-cells, which recognise self-lipids presented by the antigen-presenting molecule CD1c, are frequent in human blood, but their role during infection is unclear. Here, we investigate how CD1c-expressing antigen-presenting cells (APCs) and Mtb infection shape CD1c-autoreactive T-cell responses using engineered human APC systems, complemented by single-cell transcriptomic profiling to define the *ex vivo* phenotypic landscape of these T-cells. CD1c is present within human TB granulomas, whereas Mtb down-modulates CD1c expression on infected APCs, consistent with an immune evasion strategy. CD1c-autoreactive T-cells respond more strongly to Mtb-infected CD1c+ APCs than to uninfected cells, exhibiting enhanced activation, cytotoxicity, and diverse cytokine secretion via CD1c-dependent recognition. Under *in vitro* conditions, these T-cells reduce relative Mtb burden in infected phagocytes. Single-cell RNA-sequencing reveals cytotoxic effector-memory programmes and expression of antimicrobial molecules, providing a mechanistic basis for these responses. Together, these findings define a human CD1c-restricted T-cell response to Mtb-infected APCs and identify autoreactive CD1c-restricted T-cells as a candidate cellular axis for lipid-directed immunity in TB.

## Introduction

Tuberculosis (TB), caused by *Mycobacterium tuberculosis* (Mtb), is a major global epidemic ^1–3^. The only licensed vaccine, Bacille Calmette–Guérin (BCG), has variable efficacy against adult pulmonary disease, which drives transmission, and therefore new vaccines are urgently needed ^4^. Typically, novel TB vaccines have focussed on eliciting conventional T-cell responses to peptide antigens presented by highly polymorphic major histocompatibility complex (MHC) molecules ^5,6^. Although progress has been made in developing new immunisation tools, one of the most promising candidates, M72, based on immunogenic peptides, achieved only 50% efficacy against TB ^7–10^. This partial efficacy highlights the need to explore alternative approaches, including non-peptide antigens, to achieve the level of protection required for global TB control. Consequently, there is growing interest in harnessing unconventional T-cell responses as a complementary or alternative strategy ^11,12^. Generally, unconventional T-cells recognise antigens bound to non-polymorphic antigen presenting molecules and therefore are not restricted by genetic variability. This group includes MR1-restricted mucosal associated invariant T-cells (MAIT), CD1d-restricted invariant natural killer T-cells (iNKT), HLA-E- and CD1 group 1-restricted T-cells ^11,13^.

The CD1 family (CD1a, CD1b, CD1c, CD1d) are non-polymorphic MHC-class I-like proteins that bind and present self or foreign lipid antigens to αβ and γδ T-cells ^14^. CD1c is the most ubiquitously expressed group 1 CD1 molecule, present on antigen presenting cells (APCs) such as dendritic cells (DCs), foamy macrophages and B-cells ^15,16^. Several studies suggest a role for CD1c in host immunity to Mtb. CD1c presents Mtb-derived lipids to T-cells, including mannosyl-β1-phosphomycoketide (MPM) and phosphomycoketide (PM) ^17–21^. CD1c-restricted and Mtb lipid-specific T-cells expand in the circulation of TB patients, and CD1c-PM loaded tetramers detect PM-specific T-cells in individuals with latent TB ^18,22^.

However, it is notable that the majority of CD1c-restricted T-cells exhibit autoreactivity against CD1c expressing APCs in the absence of exogenous microbial antigens, suggesting recognition of CD1c bound to endogenous self-lipids ^23,24^. These CD1c-autoreactive T-cells recognise diverse self-lipids including phospholipids, cholesteryl-esters, methyl-lysophosphatidic acids (mLPAs), and monoacylglycerols (MAG) ^25–27^. CD1c-autoreactive T-cells are relatively abundant in the circulation of healthy adults and are proposed to become activated in response to host lipids in autoimmune disease and cancer ^23^. CD1c-autoreactive T-cells isolated from patients with systemic lupus erythematosus (SLE) provide help to CD1c^+^ B-cells, enhancing the production of IgG and contributing to immunopathology ^28^. CD1c-autoreactive T-cells infiltrate lesions in Hashimoto’s thyroiditis and Graves’ disease, contributing to tissue destruction ^29^. CD1c-autoreactive T-cells recognise tumour associated self-lipid antigens (mLPA) derived from leukemic cells ^27^. mLPA-specific T-cells efficiently kill CD1c^+^ acute leukaemia cells and protect immunodeficient mice against CD1c^+^ human leukaemia cells ^27^. Therefore, CD1c-autoreactive T-cells contribute to the immune response in human autoimmune disease and tumour immunosurveillance. However, historically these diseases will not have exerted the dominant evolutionary pressure relative to mortality from infection.

Humans have co-evolved with bacteria including gut commensals ^30^ and environmental and pathogenic mycobacteria ^31^. This long-standing relationship with microorganisms may have exerted evolutionary pressure on CD1c-reactive T-cells to contribute to immune defence against pathogenic organisms ^31^. Nevertheless, although CD1c-autoreactive T-cells are well described in human blood and have been implicated in autoimmunity and cancer, their role during infection remains poorly defined. This is an important knowledge gap because CD1c is positioned to present infection-altered lipid antigens, and autoreactive CD1-restricted T-cells may respond not only to steady-state self-lipids but also to infection-induced changes in lipid antigen presentation. Indeed, CD1c-autoreactive T-cells expand after Mtb infection of CD1 transgenic mice (hCD1Tg) *in vivo* ^32^. Furthermore, autoreactive T-cells restricted by other CD1 family members also exhibit enhanced responses to TLR agonists, mycobacteria, and mycobacterial lipids, thereby indicating that these T-cells may also be activated during infection ^33,34^. However, whether human CD1c-autoreactive T-cells directly respond to Mtb-infected APCs, whether this response is CD1c-TCR-dependent, and whether these cells mediate cytotoxic or antimicrobial effector functions has not been established.

Here, we define the response of human CD1c-autoreactive T-cells to Mtb-infected APCs using complementary human *in vitro* systems. We first examine CD1c expression in human TB lung tissue and determine how Mtb infection affects CD1c expression on primary DCs and engineered THP1-CD1c APCs. We then use CD1c-expressing APCs to expand and validate CD1c-autoreactive T-cell lines, and test their activation, cytokine secretion, cytotoxicity and capacity to reduce relative Mtb burden in infected phagocytes. To establish whether enhanced recognition of infected APCs can be mediated through the TCR, we clone dominant CD1c-reactive TCRs and test their function in a Jurkat expression system. Finally, we use single-cell transcriptomic profiling of *ex vivo* CD1c-autoreactive T-cells to define their phenotypic landscape and effector programmes. Together, these data identify a previously unrecognised human CD1c-autoreactive T-cell response to Mtb-infected APCs and support these cells as candidate contributors to lipid-directed immunity during TB infection.

## Results

### Circulating CD1c-autoreactive T-cells in healthy donors

CD1c is an MHC class I-like molecule consisting of a heavy chain and β2-microglobulin (β2m) light chain26. CD1c-autoreactive T-cells are defined by their recognition of CD1c in the absence of exogenous lipid antigen ^24,25,27,35^. To investigate the presence of CD1c-autoreactive T-cells in the blood of healthy donors, we developed novel *in vitro* assays to allow their specific expansion and subsequent detection by flow cytometry. To provide a controlled CD1c-expressing APC system while minimising conventional MHC-restricted T-cell activation, we generated THP1 knockout cells lacking both β2-microglobulin (β2m) and the Class II transactivator (CIITA). Loss of β2m removes surface expression of β2m-dependent molecules, including classical MHC class I and endogenous CD1 proteins, while CIITA knockout prevents MHC class II expression. Flow cytometric analysis confirmed that WT THP1 cells expressed β2m and classical MHC class I, with low detectable MHC class II and HLA-E (Fig. 1A). In contrast, THP1-KO cells lacked detectable β2m, MHC class I, MHC class II, HLA-E, CD1b and CD1c. To provide specific CD1c stimulation, THP1-KO cells were transduced with a CD1c-β2m fusion construct to generate THP1-CD1c APCs. These cells expressed robust surface CD1c without detectable classical and non-classical MHC molecules (Fig. 1A). Thus, the engineered THP1-CD1c system provides CD1c expression in the absence of detectable surface MHC-I or MHC-II, supporting its use to assess CD1c-mediated T-cell responses.

**Fig. 1.**
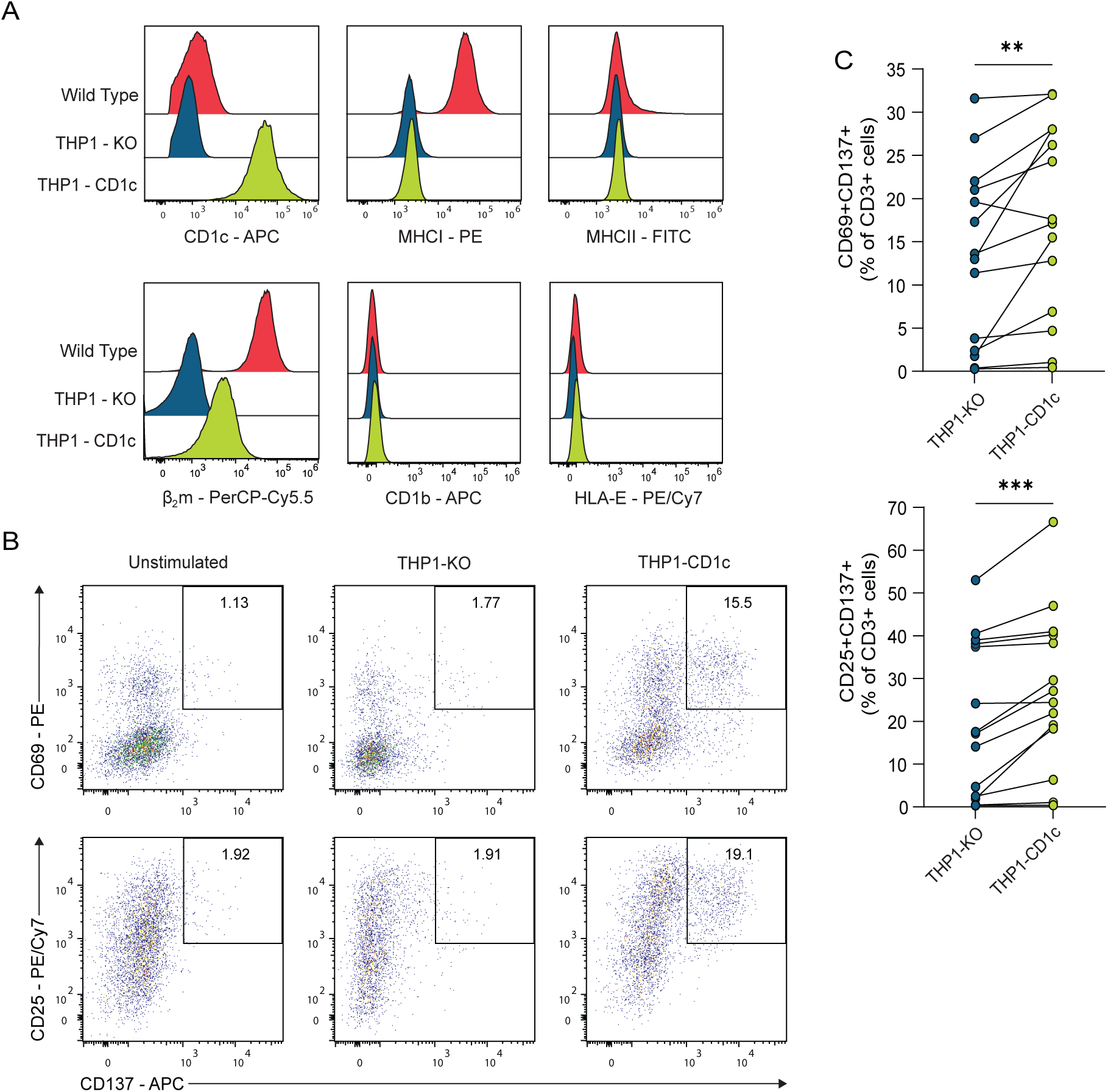
Autoreactive T-cells expand upon CD1c stimulation. (A) Representative histogram overlays showing CD1c, MHCI, MHCII, β2m, CD1b and HLA-E expression on wild type THP1 cells, and engineered THP1-KO and THP1-CD1c APCs. (B) Representative flow cytometry analysis of T-cells from a healthy donor expanded with THP1-CD1c cells in the absence of exogenous lipid antigen and then cultured overnight with THP1 cells. Cells were gated on live CD3^+^CTV^-^ T-cells and T-cell activation was measured with anti-CD69, anti-CD25 and anti-CD137. Plots show expression of CD69 and CD137 (top), CD25 and CD137 (bottom). Significant numbers of CD1c-autoreactive T-cells were present in expanded cultures. (C) Cumulative data from 14 healthy donors showing frequency of CD3^+^CTV^-^CD69^+^CD137^+^ T-cells (top), and CD3^+^CTV^-^CD25^+^CD137^+^ T-cells (bottom) from lines first expanded with THP1-CD1c and then overnight culture with THP1-KO or THP1-CD1c APCs. CD1c-autoreactive T-cells were present in the majority of donors. ** P < 0.01; *** P < 0.001; (C, Wilcoxon matched pairs signed rank test).

Purified CD3^+^ T-cells isolated from 14 healthy donors were labelled with the proliferation marker CellTrace violet (CTV) and co-cultured with irradiated THP1-CD1c APCs without exogenously added lipid antigens. After 12 days *in vitro* culture, a fraction of cells had undergone cell division, as indicated by dilution of CTV, consistent with the presence of autoreactive T-cells (Fig. S1A). To confirm this, we performed an activation-induced marker (AIM) assay and observed that a significant fraction of proliferating cells expressed the activation-induced markers CD69, CD25 and CD137 upon re-challenge with THP1-CD1c cells (Fig. 1B and 1C), with donor-to-donor variation in expansion. Importantly, CD1c-autoreactive T-cells expanded in most donors, with increased expression of CD69, CD25 and CD137 after overnight stimulation with THP1-CD1c cells (Fig. 1C). The absence of activation in response to THP1-KO cells, together with the lack of detectable MHC-I and MHC-II on THP1-CD1c APCs, supports CD1c-dependent rather than conventional MHC-restricted activation. We performed a Luminex assay on short term cultures expanded by THP1-CD1c APCs to measure cytokine secretion, studying proinflammatory and anti-inflammatory cytokines known to be critical in anti-mycobacterial immunity5. CD1c-mediated stimulation, without exogenously added antigen, induced production of several cytokines including IL-1β, IL-2, IL-10, IL-13 and GM-CSF, suggesting a polyfunctional phenotype (Fig. S1B). Thus, CD1c-autoreactive T-cells can be detected in healthy donors and expanded in short-term *in vitro* culture.

### CD1c expression in human TB granulomas

We hypothesised that CD1c-autoreactive T-cells may have a role in mycobacterial infection, and so first investigated CD1c expression at the site of TB infection, which has not been previously characterised. We performed immunohistochemical staining of lung biopsies taken from patients with active pulmonary TB with a recombinant anti-CD1c antibody. CD1c expression was observed across lung biopsies from all five TB patients, although staining was spatially heterogeneous rather than uniformly distributed throughout granulomatous lesions (Fig. 2A). Panels 2Ai–iii show granulomatous regions, including central granuloma areas where CD1c staining was infrequent. In contrast, stronger CD1c staining was observed in distal inflammatory tissue remote from the granuloma centre and in lymphoid/B-cell follicle-rich regions (Fig. 2Av-2Avi). Therefore, these data confirm the presence of CD1c-expressing cells in human TB lung tissue, while indicating that CD1c expression is spatially restricted and most prominent away from the granuloma centre.

**Fig. 2.**
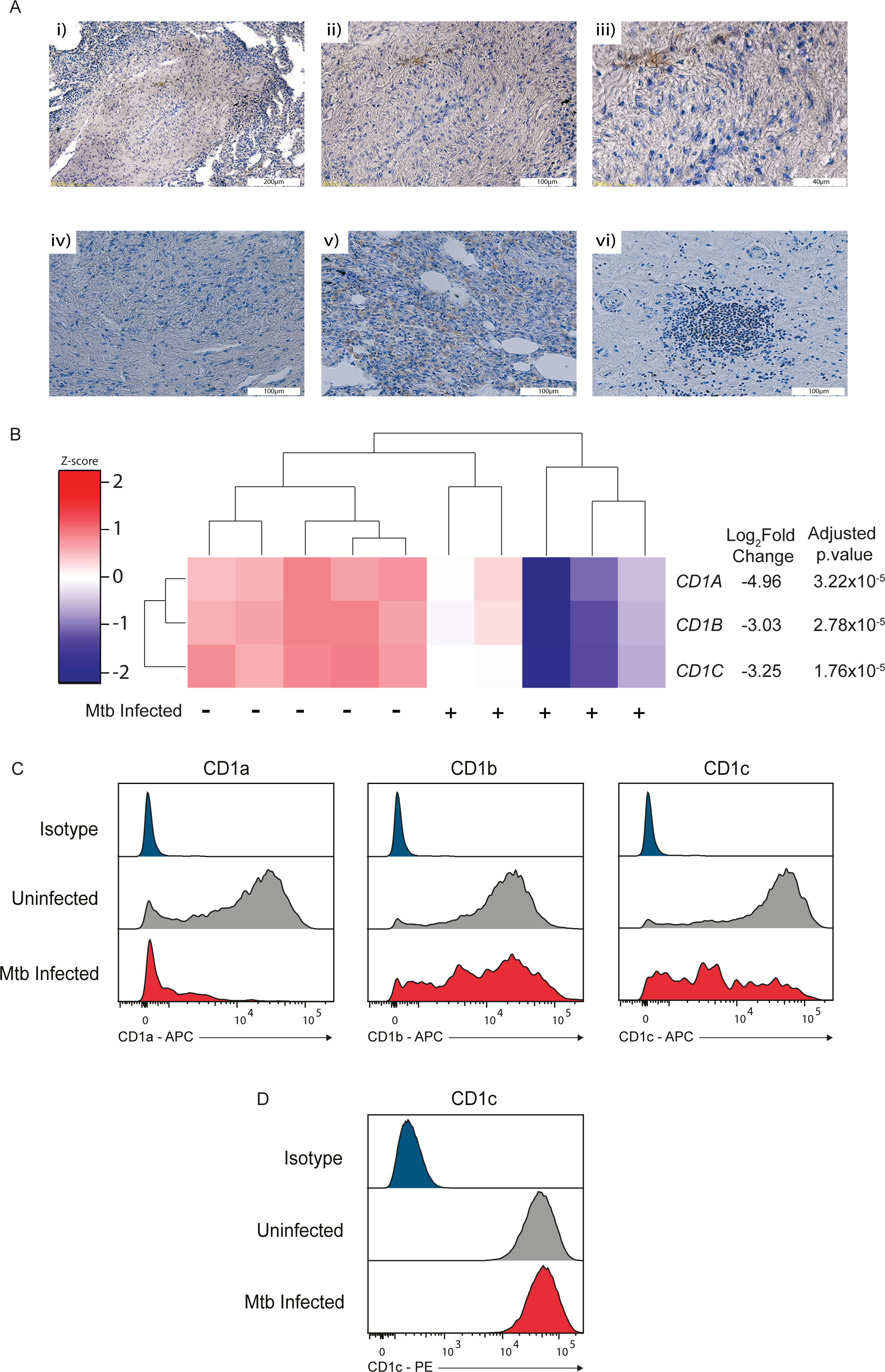
CD1c is expressed in human TB and downregulated in APCs by Mtb infection. (A) Lung biopsies from patients with active TB were stained with anti-CD1c antibodies. i-iii) Representative granulomatous regions, including central granuloma areas where CD1c immunoreactivity is infrequent. iv) Control staining of the same granuloma with secondary antibody only and avidin biotin-peroxidase complex (ABC) detection shows no immunoreactivity. v) CD1c staining is more apparent in inflammatory tissue remote from the TB granuloma centre. vi) CD1c staining in B-cell follicles adjacent to the granuloma. (B) RNA-Seq heat map showing significant reduction in CD1A, CD1B, and CD1C expression in MoDCs from five donors at 48 hours after Mtb infection (MOI=1). Differential gene expression was performed on filtered normalised counts using the voom – limma pipeline in R Studio with p-value adjustment being performed using the Benjamini-Hochberg method. Differentially expressed genes were identified as having Log2FC > +1 (Upregulated) or < -1 (Downregulated) with adjusted p-values <0.05. (C) MoDC histograms of normalised counts demonstrating a reduction in CD1a, CD1b, and CD1c expression on differentiated MoDCs following live Mtb infection (MOI=1). Data representative of an experiment conducted in two donors performed in triplicate. (D) Flow cytometry histograms showing CD1c expression on THP1-CD1c cells at 72 hours following infection with live Mtb (MOI=1). Mtb infection does not change CD1c expression on THP1-CD1c cells. Data representative of two experiments performed in triplicate.

Given the relatively low frequency of CD1c staining in central granuloma regions, we next investigated the dynamics of CD1c expression during Mtb infection. Previous reports suggested that Mtb regulates the antigen presenting functions of DCs by downregulating CD1c expression, thereby repressing their response to infection ^36,37^. To investigate the impact of Mtb infection on CD1c expression, we first interrogated an RNA-sequencing dataset derived from five healthy donors where monocyte derived DC (MoDC) were infected with live Mtb ^38^. At 48 hours after Mtb infection, CD1 group 1 expression was significantly reduced in infected MoDCs derived from all five donors relative to uninfected cells (Fig. 2B). Next, we validated these results at the protein level on MoDCs derived from healthy donors by flow cytometry. Our results corroborate the transcriptomic dataset and reveal significantly reduced expression of CD1c protein on primary MoDCs after 48 hours of live Mtb infection measured by flow cytometry (Fig. 2C & Fig. S2A). To investigate whether Mtb regulates CD1c expression on our engineered THP1-CD1c APCs, we measured CD1c expression after Mtb infection. Contrary to the reduced expression of CD1c on infected primary MoDCs, CD1c expression on THP1-CD1c APCs did not change following Mtb infection (Fig. 2D & Fig. S2B). This likely reflects the engineered nature of the THP1-CD1c system, in which CD1c is expressed from a CD1c-β2m fusion construct rather than the endogenous CD1C locus. Thus, this model allows CD1c-dependent T-cell responses to Mtb-infected APCs to be assessed without the confounding effect of infection-induced CD1c loss. This was important for subsequent functional experiments, because primary DCs infected with Mtb would be expected to lose endogenous CD1c expression, making it difficult to distinguish reduced T-cell activation caused by CD1c downregulation from changes in CD1c-presented antigens. Overall, our results reveal spatially heterogeneous CD1c expression in human TB lung tissue, with staining most apparent away from the granuloma centre, and show that Mtb downregulates CD1c on primary DCs but not in the engineered THP1-CD1c APC model used for functional assays.

### CD1c-autoreactive T-cells are cytotoxic

To investigate CD1c-autoreactive T-cell function in more detail, we derived highly enriched CD1c-autoreactive T-cell lines from the peripheral blood of healthy donors using two independent approaches. First, we expanded T-cells from a donor that exhibited robust proliferation in response to THP1-CD1c APCs (Fig. 1B), followed by CD1c-endo tetramer-guided flow cytometric sorting and *in vitro* expansion. In parallel, we enriched blood-derived T-cells from an independent donor using CD1c-endo streptamers, followed by CD1c-endo dextramer sorting and expansion (Fig. S4). The gating strategy, sorting approach and post-expansion validation are shown in Fig. S4. After expansion, sorted cells showed strong staining with CD1c-endo tetramers, whereas unstained cells and an irrelevant tetramer showed no detectable staining, confirming specific enrichment of CD1c-endo-binding T-cells. The resulting donor-derived T-cell lines showed near-uniform CD1c-endo tetramer staining after expansion, consistent with highly efficient enrichment of CD1c-endo-binding T-cells (Fig. 3A and 3C). Both lines expressed αβ TCR and CD4, with no detectable γδ TCR or CD8 expression (Fig. 3B and 3D). Functionally, both lines were activated in response to THP1-CD1c APCs but not parental THP1-KO APCs, as shown by upregulation of CD69 and CD25 (Fig. 3E). Together, these data demonstrate that these T-cell lines are highly enriched, CD1c-endo tetramer-positive, αβTCR+CD4+ T-cell populations with CD1c-dependent functional reactivity.

**Fig. 3.**
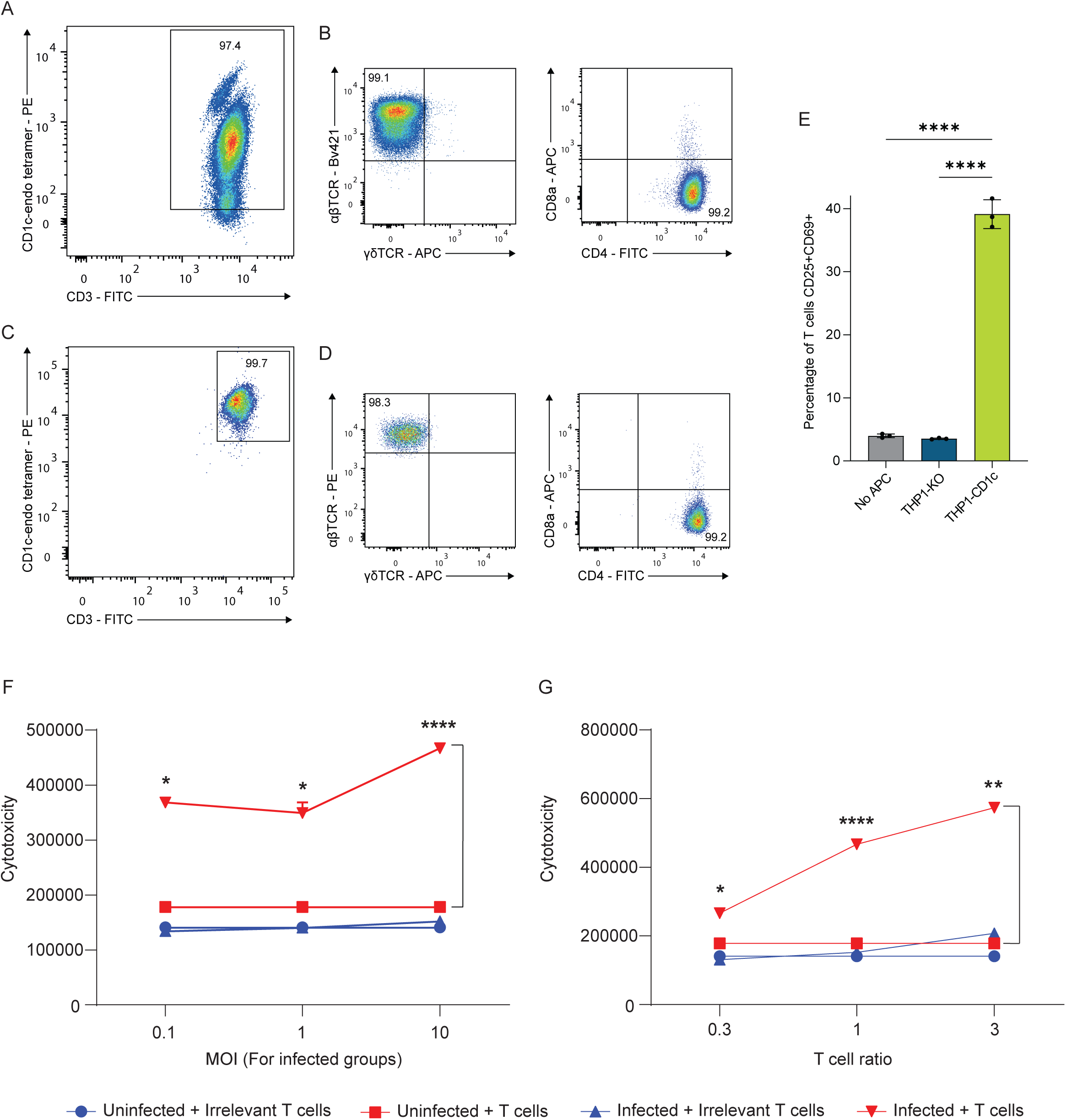
CD1c-autoreactive T-cells are cytotoxic to Mtb stimulated cells. (A-D) CD1c-endo tetramer staining of T-cell lines generated from two healthy donors. (A) T-cells generated from PBMCs after CD1c-endo streptamer enrichment and one round of CD1c-endo dextramer flow cytometry sorting and subsequent *in vitro* expansion (Fig. S3). (C) T-cells generated after expansion with THP1-CD1c cells, followed by CD1c-endo-tetramer guided cell sorting and subsequent *in vitro* expansion. (B) and (D) FACS plots demonstrating that T-cells in (A) and (C), respectively, are αβ TCR and CD4 positive. (E) T-cell lines are activated in response to THP1-CD1c APCs. Rested T-cells and T-cells cultured with THP1-KO APCs served as control. T-cell activity was determined by measuring the upregulation of CD69 and CD25 by flow cytometry. Data is representative of three experiments from the two donor-derived lines performed in triplicate. (F) CD1c-autoreactive T-cells display significant cytotoxicity in a dose-dependent manner against THP1-CD1c APCs treated with increasing doses of UV-killed Mtb. The untreated/no-infection condition represents MOI = 0. The indicated MOI values refer only to THP1-CD1c APCs treated with UV-killed Mtb. No cytotoxicity was observed by T-cells cultured with untreated THP1-CD1c APCs or irrelevant control T-cells, lacking CD1c restriction. (G) CD1c-autoreactive T-cells displayed significant cytotoxicity in a T-cell dose-dependent manner against UV-killed Mtb-treated THP1-CD1c APCs. No cytotoxicity was observed by T-cells cultured with untreated THP1-CD1c APCs or with irrelevant control T-cells. Cytotoxicity was measured using a ToxiLight assay. Data is representative of two independent experiments, each preformed in triplicate. * P < 0.05; ** P < 0.01; **** P < 0.0001 (E, one-way ANOVA with Tukey’s multiple comparison test; F-G, two-way ANOVA).

CD1c-autoreactive T-cells have been reported as having cytotoxic activity 27,39, and so we next investigated their cytotoxicity against target cells stimulated with UV-killed Mtb. We first cultured CD1c-autoreactive T-cells with THP1-CD1c APCs treated with increasing doses of UV-killed Mtb and measured cell death. CD1c-autoreactive T-cells mediated cytotoxicity toward THP1-CD1c APCs treated with UV-killed Mtb, in a Mtb dose dependent manner, while no cytotoxicity occurred against untreated THP1-CD1c APCs (Fig. 3F). Irrelevant control T-cells did not induce notable cytotoxicity against THP1-CD1c APCs (Fig. 3F). In addition, CD1c-autoreactive T-cells induced significant cytotoxicity, in a T-cell dose dependent manner, against THP1-CD1c APCs treated with UV-killed Mtb (MOI=10), but not to untreated controls (Fig. 3G). Increasing the dose of the control T-cells did not induce cytotoxicity (Fig. 3G). These results demonstrate that CD1c-autoreactive T-cells are cytotoxic towards CD1c^+^ APCs treated with UV-killed Mtb.

### CD1c-autoreactive T-cells exhibit enhanced lysis of Mtb-infected APCs beyond baseline autoreactivity

To study T-cell cytotoxicity in more detail, we performed complementary assays studying live Mtb infection. We cultured T-cells with THP1-KO and THP1-CD1c that were either uninfected or infected with live Mtb. First, we investigated T-cell activity by measuring the activation markers CD69 and CD25 by flow cytometry. CD1c-autoreactive T-cells were activated when cultured with uninfected THP1-CD1c APCs but not with control THP1-KO APCs, consistent with CD1c-dependent autoreactivity (Fig. 4A and 4B). Notably, increased activation of CD1c-autoreactive T-cells occurred when they were cultured with Mtb-infected THP1-CD1c APCs (Fig. 4A and 4B). To investigate T-cell cytotoxicity more directly, we cultured the T-cells with APCs before measuring target cell viability by flow cytometry (Fig. 4C). Importantly, we observed lysis of Mtb-infected THP1-CD1c APCs by CD1c-autoreactive T-cells, greater than the lysis of uninfected THP1-CD1c APCs (Fig. 4D).

**Fig. 4.**
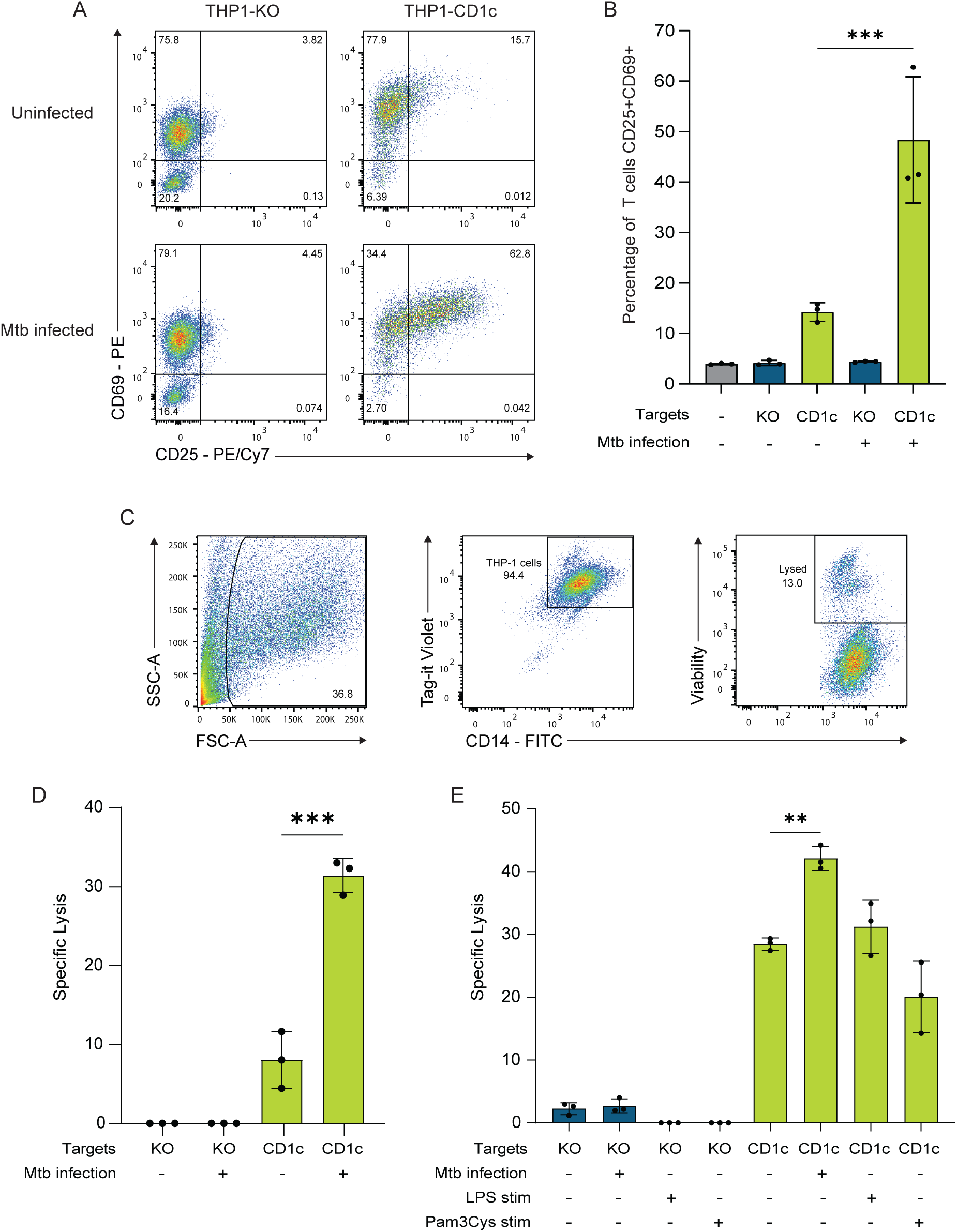
CD1c-autoreactive T-cells lyse target cells infected with live Mtb. (A) Flow cytometry dot plots and (B) bar graphs showing CD1c-autoreactive T-cells are activated by THP1-CD1c APCs but not THP1-KO APCs, with significantly greater activation when THP1-CD1c APCs are infected with live Mtb (MOI=1). (C) Gating strategy for measuring the T-cell mediated lysis of THP1 cells. Prior to T-cell culture, THP1 cells were stained with Tag-it violet to allow identification of THP1 target cells in co-culture. LIVE/DEAD Fixable Near-IR Dead Cell Marker was used to measure the proportion of dead THP1 cells. Matched APC-only controls, including uninfected and Mtb-infected THP1 cells cultured without T cells, were included in parallel to define baseline target-cell death for each condition. (D) CD1c-autoreactive T-cells lyse THP1-CD1c APCs but not THP1-KO APCs. Killing is significantly enhanced when THP1-CD1c cells are infected with live Mtb (MOI=1). Specific lysis was calculated by subtracting baseline target-cell death observed in matched APC-only controls from target-cell death observed in the corresponding T-cell co-culture condition. (E) Lysis of THP1-CD1c is increased with Mtb infection, but not enhanced with LPS or Pam3CSK4 stimulation. Specific lysis was calculated relative to matched APC-only controls cultured in parallel. Data are representative of three independent experiments, each performed in triplicate. ** P < 0.01; ***P < 0.001 (B, D and E, one-way ANOVA with Tukey’s multiple comparison test).

Recent studies suggest that activation of APCs with TLR agonists can enhance the activation of CD1-autoreactive T-cells ^33,40^. Therefore, to investigate whether the enhanced cytotoxicity of CD1c-autoreactive T-cells was facilitated via TLR mediated mechanisms, we cultured T-cells with THP1-CD1c APCs that were infected with Mtb or stimulated with the TLR agonists Pam3Cys (TLR2) or LPS (TLR4). TLR agonists did not enhance T-cell cytotoxicity above the autoreactive response, whereas Mtb infection of APCs induced significant cytotoxicity of THP1-CD1c APCs (Fig. 4E). Overall, our results confirm that CD1c-autoreactive T-cells kill Mtb-infected CD1c^+^ APCs, which is not replicated by TLR activation alone.

### The enhanced Mtb reactivity is mediated via CD1c recognition by the TCR

To provide further mechanistic insight, we performed single-cell TCR sequencing of CD1c-autoreactive T-cells (Fig. 3A). Single CD1c-endo tetramer-positive T-cells were sorted into individual wells for targeted TCR sequencing. After filtering and manual curation, 11 single cells yielded productive paired αβ TCR sequences. The repertoire was oligoclonal, with two dominant productive clonotypes accounting for 10 of 11 paired TCRs. The dominant clonotype, designated EM1, was detected in 6 of 11 cells and consisted of an alpha chain containing TRAV4 and TRAJ5 paired with a beta chain containing TRBV7-8, TRBD1, and TRBJ2-7. The second clonotype, designated EM2, was detected in 4 of 11 cells and consisted of an alpha chain containing TRAV1-2 and TRAJ26 paired with a beta chain containing TRBV2, TRBD1, and TRBJ1-1 (Fig. 5A). We utilised the β2m knock-out JRT3.5 Jurkat T-cell line, which lacks both TCR and CD1c expression, to avoid self-reactivity by Jurkat T-cells in functional studies (Fig. S4). This reporter system was used to test whether EM1 and EM2 confer CD1c-dependent recognition and activation, rather than cytotoxic effector function. We transduced the β2m knock-out JRT3.5 Jurkat T-cell line with the EM1 and EM2 TCRs. Staining of TCR transduced Jurkats with CD1c-endo tetramers showed clear binding of EM1 and EM2 Jurkat T-cells, consistent with direct engagement of CD1c by the TCR (Fig. 5B). CD1c-endo tetramers did not stain the control parental Jurkat T-cells (Fig. 5B). In addition, culturing the Jurkat T-cells on plates coated with CD1c-endo protein resulted in activation of both EM1 and EM2 Jurkat T-cell lines demonstrated by upregulation of CD69, suggesting TCR-dependent signalling via CD1c engagement (Fig. 5C).

**Fig. 5.**
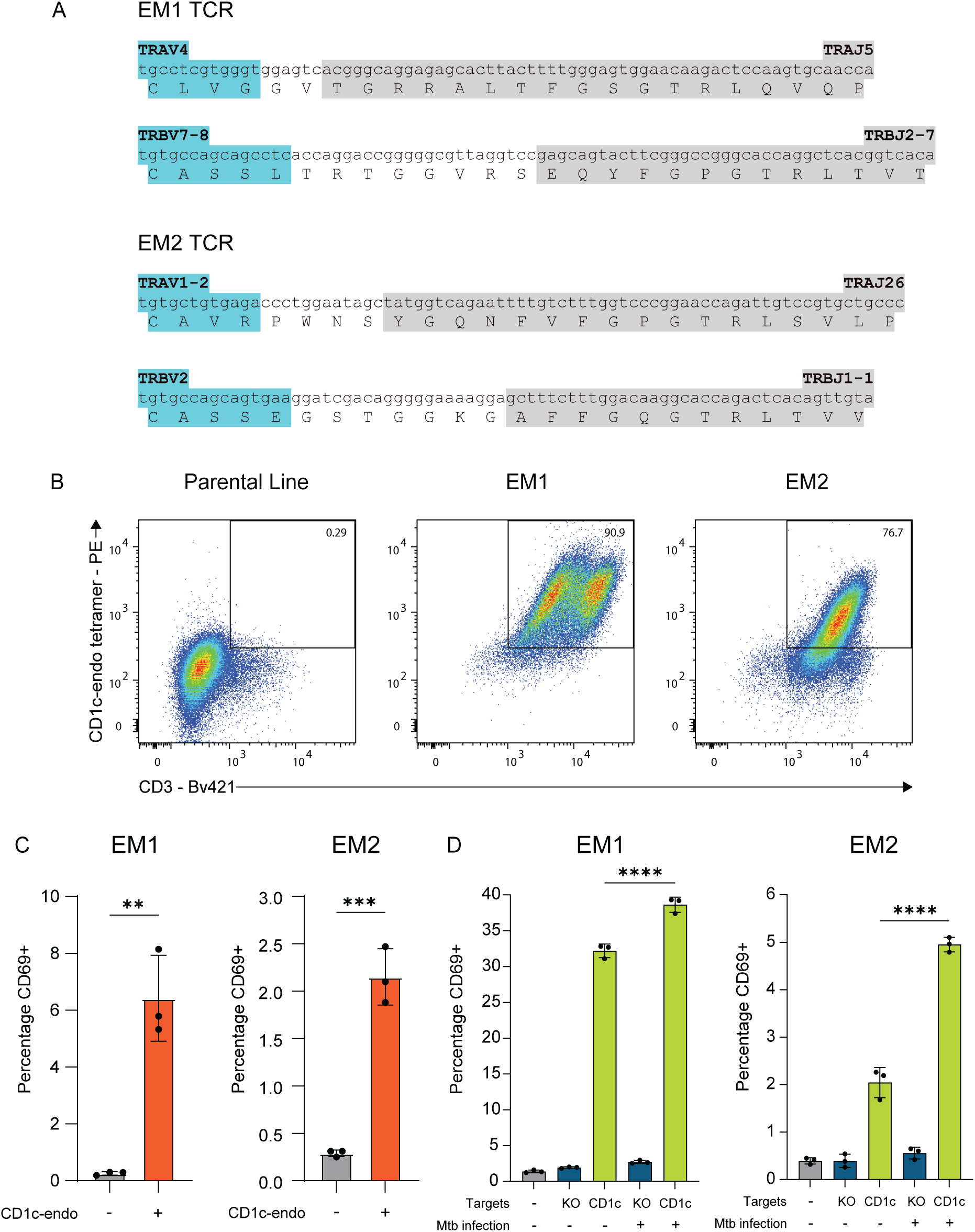
TCR-CD1c interactions mediate the response to Mtb infection. (A) The alpha (top) and the beta (bottom) chain sequences of EM1 and EM2 TCRs. Variable region (grey), N additions (white) and joining segment (blue) are shown. After filtering and manual curation, 11 single cells yielded productive paired αβ TCR sequences. EM1 and EM2 represented 6/11 cells and 4/11 cells, respectively. (B) CD1c-endo tetramer staining of parental Jurkats and Jurkat T-cells transduced to express EM1 and EM2 TCRs. (C) Percentage activation of Jurkat T-cells transduced with EM1 and EM2 TCRs in response to uncoated or CD1c-endo coated wells. T-cell activity was measured by flow cytometry staining with anti-CD69. (D) Jurkat T-cells transduced with EM1 and EM2 TCRs are activated by THP1-CD1c APCs but not THP1-KO APCs, with significantly greater activation when THP1-CD1c APCs are infected with live Mtb (MOI=1). Data are representative of two independent experiments, each performed in triplicate. ** P < 0.01, *** P < 0.001, **** P < 0.0001 (C, unpaired t-test, D, one-way ANOVA).

Mtb infection downregulates endogenous CD1c on primary MoDCs (Fig. 2B and 2C), and therefore we tested EM1 and EM2 TCR reactivity using the engineered THP1-CD1c system, in which CD1c expression is preserved during infection (Fig. 2D). We observed Jurkat T-cell line activation through upregulation of CD69 when cultured with uninfected THP1-CD1c APCs but not with control THP1-KO APCs, confirming the autoreactivity of the TCRs (Fig. 5D). Importantly, we observed significantly more Jurkat T-cell activation when cells were cultured with Mtb-infected THP1-CD1c APCs (Fig. 5D). These data suggest that the enhanced activation of CD1c-autoreactive T-cells against Mtb-infected target cells is mediated by TCR recognition of CD1c.

### CD1c-autoreactive T-cells release cytokines associated with TB immunity and suppress Mtb luminescence

To further investigate T-cell function in the context of Mtb infection, we cultured CD1c-autoreactive T-cells with APCs that were treated with UV-killed Mtb and analysed cytokine secretion. CD1c-autoreactive T-cell cytokine secretion was increased somewhat in response to untreated THP1-CD1c APCs, further corroborating their CD1c-autoreactivity (Fig. S5A). Critically though, CD1c-autoreactive T-cells secreted higher levels of cytokines in response to Mtb-treated THP1-CD1c APCs (Fig. 6A and 6B). No cytokines were released by an irrelevant control T-cell line (Fig. 6A), and the response was CD1c-mediated, with no upregulation in response to THP1-KO cells (Fig. S5B). To determine cytokines released by the APCs alone, we treated THP1-CD1c cells with Mtb and measured their cytokine release. Mtb-treated THP1-CD1c APCs released significant amounts of IL-8 and RANTES, but not any other cytokine (Fig. S6).

**Fig. 6.**
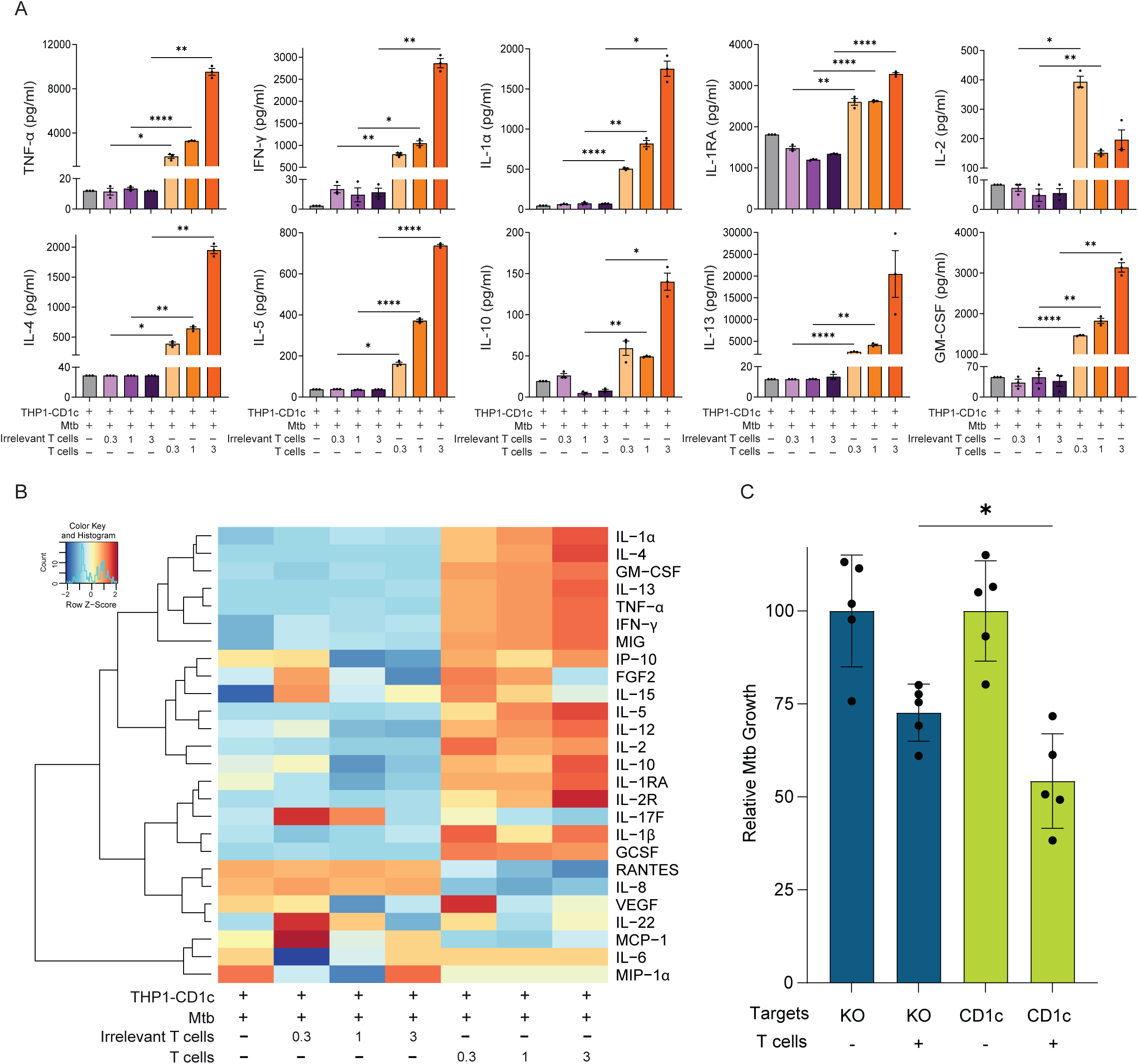
CD1c-autoreactive T-cells secrete diverse cytokines and reduce Mtb luminescence. (A) Cytokines secreted by CD1c-autoreactive T-cells cultured with Mtb-infected THP1-CD1c APCs. CD1c-autoreactive T-cells produced significant amounts of the Th1 cytokines TNF-α, IFN-γ, IL-1α and GM-CSF, and the Th2 cytokines IL-4, IL-5, IL-10 and IL-13 in a T-cell dose dependent manner. Irrelevant control T-cells did not release cytokines. (B) Heat map summarising cytokines released by CD1c-autoreactive T-cells, or irrelevant control T-cells (CD1c unrestricted), in response to Mtb-infected THP1-CD1c APCs. Red indicates high concentrations, and blue indicates low concentrations. Cytokine secretion was measured using a Luminex assay. Data are representative of two independent experiments, each preformed in triplicate. (C) CD1c-autoreactive T-cells reduce Mtb luminescence significantly when cultured with THP1-CD1c APCs relevant to when they are cultured with THP1-KO APCs. T-cells cause some reduction in THP1-KO cells, and this is greater in THP1-CD1c cells. * P < 0.05; ** P < 0.01, **** P < 0.0001 (A, Two-way ANOVA, C, unpaired t-test).

CD1c-autoreactive T-cells secreted the proinflammatory cytokines IFN-γ, TNF-α, IL-1α, IL-2 and GM-CSF in response to Mtb-treated APCs, in a dose-responsive effect (Fig. 6A and B). Additionally, CD1c-autoreactive T-cells produced IL-4, IL-5, IL-10 and IL-13, demonstrating polyfunctionality. Finally, we assessed whether CD1c-autoreactive T-cells could restrict Mtb growth in phagocytes using luminescent live Mtb. Following infection, extracellular bacteria were removed by washing before T-cell co-culture and luminescence was then monitored as a measure of viable lux-expressing Mtb. These T-cells marginally reduced Mtb growth in THP1-KO APCs, likely due to basal cytokine secretion ^41^. Importantly, luminescence was significantly further reduced in infected THP1-CD1c APCs, consistent with a CD1c-dependent reduction of relative Mtb burden, as luminescence closely correlates with colony-forming units42 (Fig. 6C). Because this assay measures luminescence of lux-expressing Mtb and does not distinguish intracellular from extracellular bacteria, we interpret these data as evidence of reduced net viable bacterial burden under *in vitro* co-culture conditions, rather than selective intracellular bacterial killing. Overall, our findings demonstrate that CD1c-autoreactive T-cell responses are significantly greater towards Mtb-infected APCs than the basal autoreactive response and reduce relative Mtb burden under these *in vitro* conditions.

### CD1c-autoreactive T-cells exhibit cytotoxic effector memory phenotype

Given their ability to restrict Mtb growth and secrete proinflammatory cytokines in response to infected APCs, we performed unbiased transcriptomic profiling to gain deeper insight into the functional properties of CD1c-autoreactive T-cells. To this end, we isolated T-cells *ex vivo* from two donors using barcoded “dCODE” CD1c-endo dextramers and performed single-cell RNA sequencing. CD3⁺ T-cells were enriched from PBMCs, and dextramer-positive and -negative T-cells were sorted for comparative analysis (Fig. 7A and Fig. S7). UMAP visualisation of the combined dataset revealed diverse T-cell populations, including naïve and effector memory (TEM) CD4⁺ and CD8⁺ T-cells (Fig. 7B and Fig. S8).

**Fig. 7.**
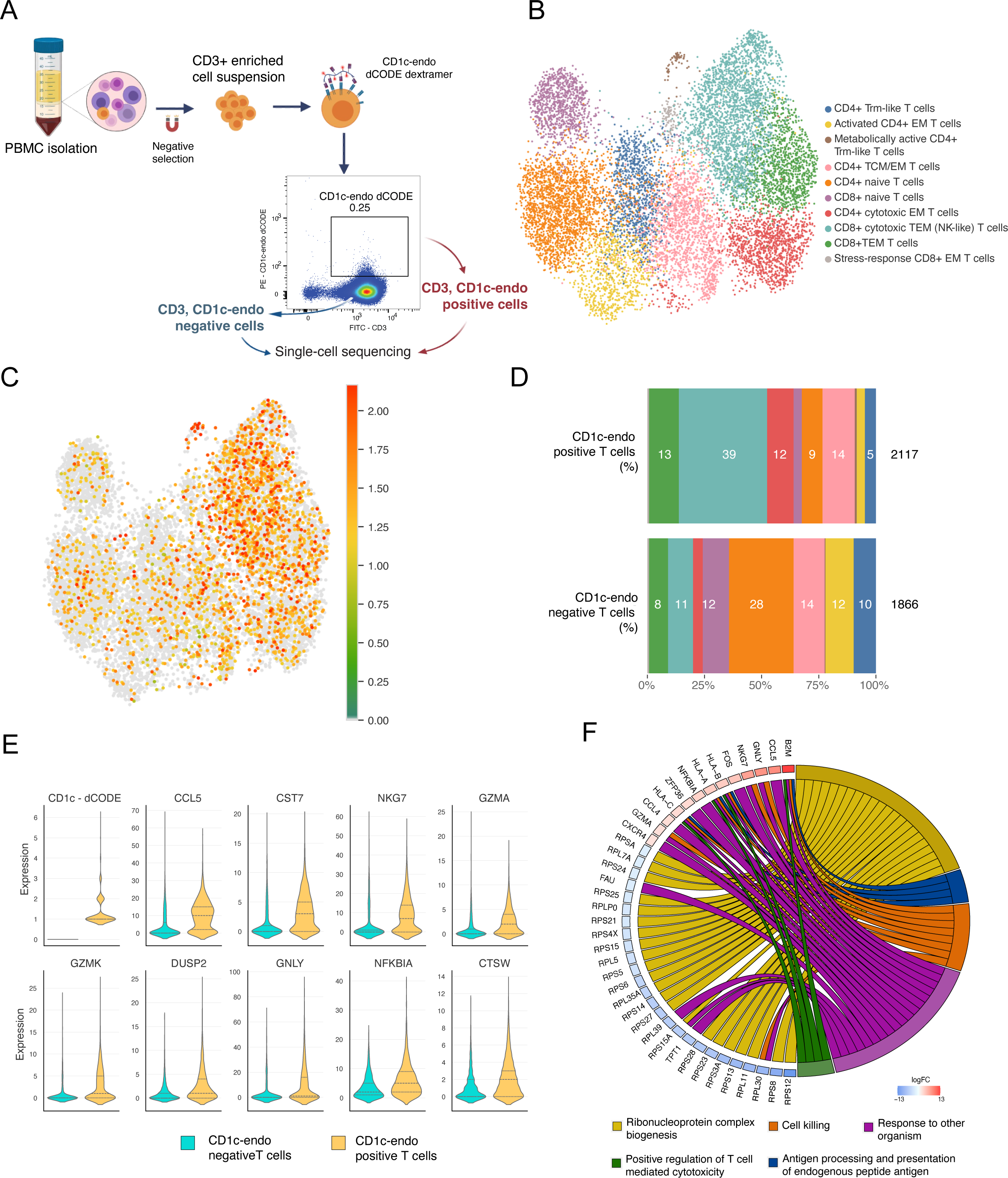
Single-cell profiling reveals that CD1c-endo dextramer positive T-cells are enriched for cytotoxic and effector phenotypes. (A) Schematic overview of the sample processing workflow. PBMCs were isolated from two donors and CD3^+^ T-cells were subsequently enriched by negative selection. Cells were stained with CD1c-endo dCODE dextramer, and CD3^+^CD1c-endo^+^ (positive) and CD3^+^CD1c-endo^−^ (negative) T-cells were sorted for single-cell RNA sequencing. (B) UMAP visualisation of 11,804 single T-cells clustered by transcriptional profile. Clusters were annotated as functional T-cell subsets, including CD4⁺ and CD8⁺ naïve, central memory (TCM), effector memory (EM and TEM), cytotoxic, and stress-response populations. Additional subsets included metabolically active T cells and tissue-resident memory-like (Trm-like) CD4⁺ cells. TRM: tissue-resident memory; EM: effector memory; TEM: T effector memory; TCM: T central memory. (C) UMAP projection showing CD1c-endo dextramer binding intensity, reflecting the number of bound dextramer molecules per cell. Cells with higher binding intensities are enriched within cytotoxic and effector CD4^+^ and CD8^+^ subsets. (D) Bar plots showing the proportional distribution of T-cell subtypes among CD1c-endo-positive (top) and - negative (bottom) populations. CD1c-endo-positive T-cells were enriched for cytotoxic and effector subsets. (E) Violin plots showing expression levels of the top 10 differentially expressed genes by Wilcoxon rank-sum test. CD1c-endo positive T-cells upregulate genes associated with cytotoxicity (e.g., NKG7, GZMA, GZMK, GNLY, CTSW) and inflammation (e.g., CCL5, NFKBIA, DUSP2), consistent with a distinct effector phenotype. (F) Chord plot illustrating functional enrichment of differentially expressed genes in CD1c-endo positive T-cells. Upregulated genes are associated with biological processes such as cell killing, antigen processing and presentation, and response to other organisms. In contrast, downregulated genes predominantly map to ribonucleoprotein complex biogenesis. The colour gradient indicates the log fold-change (logFC) in gene expression.

We focused on high-confidence CD1c-autoreactive T-cells, defined by co-positivity for CD1c-endo dextramer and dCODE barcode, and identified 2,117 such cells across both donors (Fig. S9). These cells were predominantly enriched in cytotoxic CD4⁺ and CD8⁺ effector memory clusters (Fig. 7C and 7D), although they also displayed transcriptional diversity spanning multiple T-cell states present in the broader dataset. Phenotypic analysis showed high expression of cytotoxic mediators including *GZMA*, *GZMK*, and *GNLY*, as well as effector molecules such as *CCL5* and *NKG7* (Fig. 7E). Pathway enrichment analysis revealed significant upregulation of gene sets related to response to other organisms, T-cell-mediated cytotoxicity, and cell killing (Fig. 7F). Together, these findings indicate that CD1c-autoreactive T-cells exhibit cytotoxic effector memory phenotypes and express key antimicrobial effector molecules, providing mechanistic insight into their potential effector function against Mtb-infected APCs.

## Discussion

The majority of human CD1c-restricted T-cells are activated by exposure to CD1-expressing APCs in the absence of foreign lipid antigens ^23,24^. Indeed, such CD1c-autoreactive T-cells comprise 2% of all circulating human αβ T-cells 23, and studies have suggested a functional role for these T-cells in human autoimmune disease ^29^ and cancer ^27^. However, for almost the entirety of human evolution, infection is likely to have exerted a greater selective pressure affecting survival and reproduction than autoimmune disease or cancer ^43,44^. Here, we found that polyclonal CD1c-autoreactive T-cells produced higher levels of Th1 and Th2 cytokines when stimulated with Mtb-infected APCs relative to uninfected cells, mediated increased cytotoxicity against Mtb-infected APCs in a CD1c-mediated manner, and reduced the relative intracellular Mtb burden under *in vitro* conditions42. To gain insight into the cellular programmes associated with this antimicrobial effector potential, we performed single-cell RNA sequencing of *ex vivo* isolated CD1c-autoreactive T-cells from healthy donors. These cells were transcriptionally diverse but predominantly exhibited cytotoxic effector memory phenotypes. Notably, they expressed high levels of cytolytic molecules including *GZMA*, *GZMK*, and *GNLY*, as well as effector mediators such as *CCL5* and *NKG7*, consistent with a role in direct target cell killing ^45^. Granulysin is one of the few T-cell-derived molecules with demonstrated antimicrobial activity against Mtb 45, yet few human T-cell subsets express it in this context. Combined with our finding that CD1c-autoreactive T-cells can reduce Mtb luminescence *in vitro*, this places them among a rare population with antimicrobial activity in this experimental setting. These findings provide a mechanistic explanation for how CD1c-autoreactive T-cells could mediate cytotoxicity and suppress intracellular Mtb, supporting their classification as functionally distinct effector T-cells rather than merely autoreactive bystanders. To our knowledge, this is the first single-cell transcriptional analysis of human CD1c-autoreactive T-cells, offering preliminary insight into their phenotype and potential functional diversity. Our data suggest that CD1c-responsive T-cells, previously described as “autoreactive,” respond more robustly to Mtb-infected APCs, raising the possibility that, in addition to roles in autoimmunity and cancer, these cells may also contribute to antimicrobial responses. Together, these findings address a previously unresolved question: whether human CD1c-autoreactive T-cells, known mainly in the context of autoimmunity and cancer, can functionally respond to Mtb-infected APCs. By integrating engineered APC systems, TCR transfer, cytotoxicity assays and ex vivo single-cell transcriptomics, we show that these cells exhibit CD1c-dependent activation, cytotoxicity, cytokine production and antimicrobial activity in response to Mtb-infected APCs. This links CD1c-autoreactivity to infection-associated effector function and supports these cells as candidate contributors to lipid-directed immunity in TB.

Two previous studies utilising either *in vitro* activation-based assays or CD1c-endo tetramers have confirmed the presence of CD1c-autoreactive T-cells in human peripheral blood ^23,46^. Although there was a discrepancy in their estimation of *ex vivo* frequencies for CD1c-autoreactive T-cells, both studies showed that these T-cells are a natural component of the human T-cell pool ^23,46^. Moreover, both showed that CD1c-autoreactive T-cells were enriched for CD4^+^ or DN T-cells, and exhibited memory and adaptive T-cell features closely aligned with conventional MHC-restricted T-cells ^23,46^. CD1c-autoreactive T-cells expressed diverse TCRs, including αβ, γδ and δ/αβ ^46^. Although the TCR repertoire was generally diverse, they were enriched for the TRAV17, TRAV38-1 and TRBV4-1 gene segments ^46,47^. Functionally, CD1c-autoreactive T-cell clones exhibit a Th1- or a Th0-like phenotype, releasing IFN-γ, TNF-α, GM-CSF, IL-4, and IL-5 ^23^. Corroborating these earlier studies, we detected CD1c-autoreactive T-cells in the blood of healthy donors, both αβ and γδ, and demonstrated that CD1c-autoreactive T-cells were polyfunctional. The variable expansion observed across donors likely reflects heterogeneity in CD1c-autoreactive precursor frequency, TCR repertoire composition and activation state, as expected for primary human autoreactive T-cell populations ^23^. The functional and single-cell datasets presented capture different levels of this repertoire. The functional assays used two donor-derived CD1c-autoreactive T-cell lines that were CD4+αβTCR+, whereas *ex vivo* single-cell RNA-seq revealed broader CD1c-endo-binding populations, including both CD4+ and CD8+ cytotoxic effector-memory T-cells. These approaches are therefore complementary: the T-cell lines provide mechanistic evidence of CD1c-dependent function, while single-cell profiling captures the wider phenotypic diversity of the CD1c-autoreactive compartment.

We generated CD1c-restricted αβ T-cell lines with a baseline autoreactive response that was significantly greater when responding to Mtb-infected cells, with increased activation markers, cytotoxicity and cytokine secretion, and an associated reduction in Mtb luminescence in infected APCs. The Jurkat T-cell experiments with EM1 and EM2 TCRs support a key role for the TCR in mediating response through direct recognition of CD1c. Because Jurkat cells are not cytotoxic effector cells 48, these experiments were designed to test TCR-dependent recognition and activation rather than target-cell killing. These findings suggest that changes in CD1c expression or cytokine secretion alone are unlikely to explain the enhanced T-cell activity observed towards Mtb-infected cells and instead point to a role for lipid antigen presentation on CD1c. Our findings align with a study in 2005 where autoreactive CD1 group 1 restricted T-cell clones were generated by limited dilution, after stimulation of CD4-depleted T-cells with MoDCs pulsed with hydrophobic microbial extracts derived from Mtb, *E-coli*, and *Y. enterocolitica* ^49^. A panel of 15 autoreactive clones were generated with the majority being CD1a- and CD1b-restricted and only two clones being CD1c-autoreactive ^49^. While the clones exhibited weak autoreactivity, the response to microbial antigens far exceeded the self-reactive response seen in assays of T-cell proliferation 49, similar to our findings and suggesting an antimicrobial role. TCR transfer experiments supported a role for the TCRs in promoting this dual reactivity to self and foreign antigen. However, further data from clone Mt2.33, which was a CD1c-autoreactive clone generated through Mtb antigen stimulation, was not provided. Further evidence for CD1c-restricted dual reactivity comes from a study identifying Vδ1⁺ γδ T-cells using CD1c-PM tetramers, where Jurkat cells transduced with these γδ TCRs exhibited spontaneous activation and stronger responses to microbial lipid antigens, again implicating TCR-driven recognition of CD1c-presented ligands ^50^. Another study investigated the CD1b-autoreactive clone HJ1, which was isolated from hCD1Tg mice, in *Listeria monocytogenes* infection ^33^. While HJ1 T-cells exhibited autoreactivity towards CD1b, responses were enhanced by treatment of APCs with *Listeria* or the TLR agonists Pam3Cys (TLR2) and LPS (TLR4) ^33^. Together, these observations suggest that some autoreactive CD1-responsive T-cells may have evolved as anti-microbial effectors, but when studied experimentally without such stimuli, a residual autoreactive phenotype is observed.

TLR activation during infection of APCs promotes the accumulation of stimulatory self-lipid antigens, contributing to the enhanced activation of iNKT cells ^40^. Indeed, treatment of APCs with LPS or bacteria induced the accumulation of stimulatory iNKT self-lipid agonists b-d-glucopyranosylceramides (b-GlcCer) ^40^. In our system, Mtb infection of THP1-CD1c APCs enhanced cytotoxicity by CD1c-autoreactive T-cells, but this effect was not reproduced by TLR2 or TLR4 agonists alone. This suggests that generic TLR2/4-mediated APC activation is insufficient to explain the enhanced response to Mtb-infected cells. Together with the TCR transfer experiments, these data point towards a CD1c-dependent signal that is increased or altered during Mtb infection. The molecular identity of this signal remains unresolved. One possibility is that Mtb infection alters the CD1c-presented lipid repertoire, either through presentation of Mtb-derived lipids 49,50, infection-induced host “stress lipids” 26, or bacterial and mammalian shared lipids similar to those described for CD1b-autoreactive T-cells ^34,51^. Alternatively, infection may alter lipid processing, trafficking or loading onto CD1c, increasing presentation of permissive self-lipids that can sit deeply within the CD1c groove and expose the CD1c surface for TCR recognition ^25^. Stimulatory lipids could include cholesteryl esters ^26^ or other infection-induced lipid species. Thus, the additional signal provided by Mtb-infected THP1-CD1c cells is likely to be a qualitative or quantitative change in CD1c-associated lipid presentation, rather than CD1c expression alone or generic APC activation, a hypothesis supported by our TCR transfer experiments.

A limitation of this study is that we did not directly identify the lipids presented by CD1c on Mtb-infected APCs, making it unclear whether the relevant ligands are host-derived, bacterial, or shared lipid species. Future studies should isolate CD1c molecules from Mtb-infected APCs and use mass spectrometry-based lipidomics to define the lipid species associated with enhanced T-cell activation ^52^. Comparing these autoreactive populations with non-autoreactive CD1c-restricted T-cells that recognise defined microbial lipids will also be key, as the two may represent complementary modes of lipid immune surveillance. Although Mtb luminescence correlates closely with CFU counts in this system 42, it remains an indirect measure of viable bacterial burden, and future studies using CFU plating or complementary bacterial quantification methods will be important to confirm the extent of CD1c-dependent growth restriction. A further limitation is that the single-cell RNA-seq analysis was performed on CD1c-endo-binding T-cells from two donors. Although subsampling analysis supported the robustness of the transcriptional patterns within this dataset, larger donor cohorts will be required to determine the consistency of cytotoxic effector-memory programmes across the broader human CD1c-autoreactive T-cell repertoire. In addition, our experiments rely on *in vitro* human cell systems, and *in vivo* validation of CD1c-restricted responses will benefit from specialised CD1-transgenic models that accommodate human CD1c biology, beyond what is possible in standard mouse systems_32,53._

Our observation of CD1c expression within human lung TB granulomas provides further evidence for a potential role for lipid-specific T-cell immunity in the host immune response to Mtb. Our findings are consistent with reports suggesting that CD1c expression is targeted by Mtb and BCG, leading to reduced expression on APCs such as DCs 36,37,54,55. A recent study identified a potential mechanism, with Mtb driving expression of the microRNA miR-381-3p and thereby suppressing CD1c ^37^. This has important implications for interpreting functional assays using primary DCs. Although primary CD1c-expressing DCs are biologically relevant APCs, Mtb-induced CD1c downregulation would make it difficult to determine whether altered T-cell activation reflects changes in the CD1c-presented lipid repertoire or simply reduced CD1c at the cell surface. For this reason, the engineered THP1-CD1c system provides a controlled reductionist model in which CD1c expression is maintained during infection, allowing CD1c-TCR-dependent activation to be tested directly. This difference between primary MoDCs and THP1-CD1c APCs likely reflects the engineered nature of the THP1-CD1c system.

Rather than regulation from the endogenous CD1C locus, CD1c is expressed from a heterologous CD1c-β2m fusion construct lacking the 3’-untranslated region where miR-381-3p can bind to disrupt translation37. Therefore, our data are consistent with the idea that CD1c-autoreactive responses may contribute to protective host immunity, and that Mtb-mediated suppression of CD1c expression on primary APCs may represent an immune evasion strategy37. Future studies comparing TB lesions with healthy lung and non-TB inflammatory lung tissue will be important to determine whether the spatial pattern of CD1c expression observed here is specific to TB or reflects broader inflammatory remodelling of lung APC populations.

Taken together, our findings suggest that T-cells previously described as autoreactive may also play a role in anti-mycobacterial immunity, and that this phenotype could have been shaped by evolutionary adaptation to mycobacterial exposure. Mtb has killed many more humans than any other single infection, and humans and Mtb may have co-evolved for 70,000 years 56,57, demonstrating the significant impact mycobacteria will have had on human immune development58. We identify functional responses to Mtb-infected cells by CD1c-autoreactive T-cells, including activation, cytotoxicity, cytokine secretion and reduction in Mtb luminescence *in vitro*, that clearly exceed the autoreactive phenotype. Together, our findings suggest that CD1c-autoreactive T-cells may contribute to immune responses against infection and warrant further investigation in the context of vaccine development.

## Methods

### Flow cytometry

The following fluorescent reagents were used: All antibodies were from Biolegend unless stated; anti-CD3-Bv510 (clone UCHT1); anti-CD3-FITC (clone UCHT1); anti-MHCI-PE (clone W6/32); anti-MHC-II-FITC (clone Tu39); anti-β2m-PerCP-Cy5.5 (clone A17082A); anti-αβTCR-Bv421 (clone IP26); anti-γδTCR-APC (clone B1); anti-CD4-FITC (clone RPA-T4); anti-CD8-APC (clone HIT8a); anti-CD14-FITC (clone M5E2); anti-CD11c-Bv421 (clone Bu15); anti-HLA-DR-PE/Cy7 (clone L243); anti-CD1a-APC (clone HI149); anti-CD1b-APC (clone SN13(K5-1B8)); anti-CD1c-APC (L161); anti-CD1c-PE (clone L161); anti-CD69-PE (clone FN50); anti-CD25-PE/Cy7 (clone BC96); anti-CD137-APC (clone 4B4-1); LIVE/DEAD Fixable Near-IR Dead Cell Marker and Cell Trace Violet (CTV) (both ThermoFisher Scientific); and Propidium iodide (Sigma). T-cells were treated with 50 nM Dasatinib (Axon) and blocked with Human TruStain FcX™ (Biolegend) for 30 minutes at 37°C, and anti-CD36 (clone 5-271, Biolegend) for 20 minutes before the addition of tetramers and staining reagents. Cells were stained for 45 minutes, extensively washed and acquired on a FACSAria Fusion, or a FACSAria II (All BD Biosciences). Data was analysed using FlowJo VX software (FlowJo LLC).

### Cell lines

The following cell lines were utilised in this study: THP1 (myelomonocytic leukemia), J.RT3-T3.5 (TCRβ-deficient T-cell leukemia) and HEK293TN (Human Embryonic Kidney). The engineered THP1-KO cells were generated using CRISPR-Cas9 technology to prevent the expression of β2m and Class II transactivator (CIITA). Following the knockout of β2m and CIITA, the cells were transduced using a lentivirus system with a plasmid containing a β2m-CD1c single chain gene construct to generate the THP1-CD1c cell line. Three days post transduction, CD1c^+^ cells were sorted by flow cytometry and maintained in culture. Jurkat, THP1 and HEK293TN cells were maintained in complete RPMI or DMEM (for HEK293TN) containing 10% Foetal Bovine Serum (Sigma), 1% Non-essential Amino Acids, 1% L-glutamax, 1% Sodium Pyruvate, 1% Penicillin/Streptomycin (all from ThermoFisher Scientific). All primary cell lines used in this study were isolated from peripheral blood mononuclear cells (PBMCs) obtained using Ficoll-Paque (Cytiva) from healthy blood bank donors and maintained in culture in T-cell media; RPMI containing 5% Human AB Serum (Merck), 1% Penicillin/Streptomycin, 1% L-glutamax, 1% Non-essential Amino Acids, 1% Sodium Pyruvate Pyruvate (All ThermoFisher Scientific). Pan T-cells from healthy donors purified by EasySep Human T-cell Isolation Kit (Stem cell Technologies) were labelled with CTV according to manufacturer instructions and stimulated with irradiated (80 Grays) THP1-KO or THP1-CD1c cells at a 6:1 ratio, or without THP1 cells as control. THP1 cells were irradiated to prevent APC proliferation during the 12-day T-cell expansion culture while preserving their use as stimulatory cells. On days 4, 6, 8, 10 and 12 post stimulation, 10 IU/ml rhIL-2 (Proleukin, Chiron) was added. At 12 days post stimulation, cells were rechallenged with THP1-KO or THP1-CD1c APCs overnight before staining for activation marker expression on live CD3^+^ CTV low cells by flow cytometry. Some cultures were stained with CD1c-endo tetramers directly on day 12 and then tetramer positive T-cells were sorted by flow cytometry and expanded in the presence of phytohemagglutinin (PHA) (ThermoFisher Scientific) (1µg/ml), rhIL-2 (100U/ml), and irradiated PBMC (5×105 cells/ml).

### Tetramers

Human CD1c was produced as a single chain construct with β2m N-terminally fused to the heavy chain via a flexible glycine-serine linker. Expression cassettes were cloned in-frame with C-terminal Avi-tag and His6 tag in pCDNA3.1 encoding an additional BirA Ligase. Protein expression was performed using the Expi293 Expression System (ThermoFisher Scientific). Soluble CD1c proteins were purified using nickel affinity and size exclusion chromatography. For CD1c-endo tetramers, purified CD1c-endo monomers were used to generate fluorescent-labelled CD1 tetramers by conjugation to PE-streptavidin (Biolegend). To generate dextramers, the same process was repeated as above, with PE-labelled dextran backbones (Dex-PE) (Immudex).

### TCR sequencing

Single CD1c-endo tetramer-positive cells were isolated by flow cytometric sorting into individual wells of a 96-well plate. Cells were lysed, and an oligo-dT primer was annealed to the poly-A tails of mRNA before reverse transcription to generate cDNA using an approach adapted from the SMART-seq2 protocol59. During reverse transcription, adaptor sequences were incorporated into the cDNA, enabling subsequent universal PCR (polymerase chain reaction) amplification using primers targeting the adaptor regions. Targeted PCRs were then performed for each single cell to amplify TCR variable regions using reverse primers specific for TRAC, TRBC, TRGC and TRDC constant regions. PCR products were barcoded with an 8-bp DNA sequence unique to each well. PCR products were purified, and DNA fragments of 500-600 bp were size-selected. Next-generation sequencing (NGS) libraries were prepared from size-selected fragments using New England Biolabs library preparation kits and sequenced on the MiSeq platform. Sequences covering CDR3, variable and constant regions were obtained and analysed using Seven Bridges software to generate final TCR sequences. Primer sequences are provided in Table S1.

### TCR cloning and transduction

Sequences identified from tetramer-guided sorting were constructed into full length TCR, synthesised and sub-cloned into the pELNS Lentivector by Genscript. Adherent HEK 293TN cells were co-transfected with pELNS lentivector (2.5 μg) and three accessory plasmids: pCMV-VSV-G (1.5 μg), pRSV.REV (3 μg), and pMDL.pg.RRE (3 μg), to induce production of Lentiviral particles. Lentiviral particles were harvested, filtered, and used directly for transduction of β2m knock-out J.RT3-T3.5 Jurkat cells (kind gift from Pierre Vantourout, King’s college London). Transduced cells were sorted by flow cytometry on a FACSAria IIU (BD Biosciences).

### CD1c-endo streptamers

Strep-Tactin® magnetic microbeads (IBA Lifesciences) were prepared by washing and resuspending in binding buffer (PBS, 1mM EDTA, 0.5% bovine serum albumin) prior to mixing with CD1c-endo incorporating Strep•Tag® II peptide sequence and incubated at 4°C overnight. The following day, additional CD1c-endo was added and incubated for 30 minutes at 4°C to complete the preparation of the CD1c-endo streptamer. To enrich CD1c-restricted T-cells, 2x107 T-cells were resuspended in binding buffer containing 50nM dasatinib (Sigma) containing Human TruStain FcX™ (Biolegend) and incubated at room temperature for 10 minutes. CD1c-endo streptamer was then added to T-cells and incubated on ice for 20 minutes before separating the CD1c-endo streptamer bound cells (positive fraction) and unbound cells (negative fraction) by magnetic sorting. The enrichment procedure was repeated with the negative fraction. The positive fractions were next pooled and thoroughly washed.

To remove bound T-cells in the positive fraction from the CD1c-endo streptamer, the T-cells were resuspended in biotin elution buffer and incubated on ice for 10 minutes and separated using a magnet three times. The positive and negative fractions were stained and analysed by flow cytometry. CD1c-endo dextramer stained cells from the positive fraction were sorted by flow cytometry into a single well of a 96-well plate containing T-cell media and expanded with 1μg/mL PHA, 100IU/mL rhIL-2 and 5x105 irradiated PBMCs from three different donors.

### Mtb Culture

*Mycobacterium tuberculosis* H37Rv (Mtb H37Rv) was cultured in Middlebrook 7H9 medium (supplemented with 10% ADC, 0.2% glycerol and 0.02% Tween 80) (BD Biosciences) at 37°C in an incubator with shaking at 200 rpm. Bioluminescent Mtb H37Rv lux was cultured with kanamycin (25 μg/mL). Cultures at ∼1 × 108 CFU/ml Mtb (OD = 0.6) were used for all experiments.

### Use of UV-killed and live Mtb

UV-killed and live Mtb were used for distinct experimental purposes. UV-killed Mtb was used where controlled exposure to defined amounts of Mtb-antigenic stimuli was required without ongoing bacterial replication, particularly for dose-response cytotoxicity and cytokine-release assays. Live Mtb infection was used to test whether responses observed with UV-killed Mtb were also evident during infection of APCs, and to assess T-cell activation, target-cell lysis and relative bacterial burden in the context of active host-pathogen interaction.

### MoDC generation

To generate MoDC, CD14^+^ monocytes were isolated from PBMCs by negative selection using magnetic beads (Stem cell Technologies) and cultured in the presence of 25ng/ml rhGM-CSF and 20ng/ml rhIL-4 (both from Miltenyi) for 5 days.

### CD1 expression

#### MoDC

After 3 days of culturing monocytes in the presence of rhGM-CSF and rhIL-4, some wells were infected with live Mtb (MOI=1) whereas others were left uninfected. On Day 0, 3, 5 and 7, cells were collected from wells by thoroughly washing with 3mM EDTA and stained for flow cytometry analysis. Cells were acquired on FACSAria IIU (BD Biosciences) to measure CD1a, CD1b and CD1c expression.

#### THP1 cells

THP1-KO and THP1-CD1c cells were stained with anti-CD1c-PE and its isotype control (BioLegend) and Live/Dead Aqua (Invitrogen). On day 0, some wells were infected with Mtb (MOI=1) whereas others were left uninfected. On Day 3, cells were collected from wells by thoroughly washing with 3mM EDTA and stained for flow cytometry analysis. Cells were acquired on FACSAria IIU (BD Biosciences) to confirm the expression of CD1c on THP1 cells.

### Immunohistochemistry

Tissue sections (4μm thick) were dewaxed and rehydrated before inhibiting endogenous peroxidase. Heat-induced epitope retrieval was performed prior to blocking non-specific staining. Primary antibodies, 1:500 anti-CD1c recombinant rabbit monoclonal (EPR23189-196, Abcam) was then applied to the sections and incubated overnight at 4°C. Sections incubated with buffer alone were included as a negative control. Secondary antibody, goat anti-rabbit 1:800 (2B Scientific) was applied. Sections were developed with avidin-biotin peroxidase complexes (2B Scientific) and 3,3’-diaminobenzidine tetrahydrochloride (DAB; Launch Diagnostics). Sections were counterstained with Mayer’s haematoxylin, dehydrated, cleared and mounted using XTF mounting medium (CellPath). Sections were allowed to dry before subsequent imaging using an Olympus Bx51, CC12 DotSlide microscope.

### Differential Gene Expression Analysis

PRJNA478394 fastq files were downloaded from ENA using enaBrowserTools (v1.1.0), after which transcripts in each file were quantified against a human genome template (downloaded from ftp://ftp.ensembl.org/pub/release106/fasta/homo_sapiens/cdna/Homo_sapiens.GRCh38.cdna.all.fa.gz) using alignment-free software kallisto (v0.46.1). Counts were imported into R using tximport (v1.28.0) and converted to counts per million (CPM) using edgeR (v3.14.0). Genes with summed CPM values below 14 across five samples (corresponding to the number of donors) were removed from the expression matrix, which was then normalised using TMM normalisation. Differential gene expression analysis was performed using the voom-limma (limma v3.56.2) pipeline, where normalised counts were compared between (live) Mtb-infected MoDCs and uninfected MoDCs (controls) at 48 hours. Computed p-values were adjusted for multiple hypothesis testing using the Benjamini-Hochberg method, and differentially expressed genes (> +1 log_2_FC = Upregulated, < -1 log_2_FC = Downregulated with an adjusted p.value < 0.05) were subsequently identified.

### Gene expression heatmap

Counts in the filtered, normalised expression matrix were converted to CPM, then log_2_ transformed. The expression matrix was subset to retain values for *CD1A*, *CD1B* and *CD1C* for the control and (live) Mtb-infected samples at 48 hours post-infection. Coolmap function from limma was used to generate a heatmap showing the relative change in gene expression for each sample, from the average expression for each gene, using “de pattern”. Columns were clustered using “average” method.

### Activation assays

#### T-cells

T-cells were cultured with THP1-KO and THP1-CD1c cells that were either uninfected or infected with live Mtb (MOI=1) for 48 hours at 37°C. Activation status of T-cells was measured through CD69 and CD25 expression by flow cytometry on a FACSAria IIU (BD Biosciences).

#### Jurkat T-cells

TCR transduced J.RT3-T3.5 Jurkat T-cells were cultured for 20 hours at 37°C in 96-well plates coated with CD1c-endo protein monomers (10µg/mL). TCR transduced J.RT3-T3.5 Jurkat T-cells cultured alone served as negative control. TCR transduced J.RT3-T3.5 Jurkat T-cells were also cultured with THP1-KO and THP1-CD1c cells that were either uninfected or infected with live Mtb (MOI=1) for 20 hours at 37°C. Activation status of Jurkat T-cells was measured through CD69 expression by flow cytometry on a FACSAria IIU.

#### Activation Induced Marker (AIM) assay

Pan T-cells were isolated from healthy human blood using an EasySep Human T-cell Isolation Kit (Stem cell Technologies) and stained with CTV to track proliferation. Purified T-cells are stimulated with THP1-CD1c cells for 12 days as above. On day 12, T-cells were re-stimulated overnight with irradiated THP1-KO, THP1-CD1c or left unstimulated. Proliferated T-cells were determined as CTV low and activation was measured as upregulation of CD25^+^/CD137^+^ or CD69^+^/CD137^+^ on T-cells by flow cytometry.

### Cytotoxicity assays

#### ToxiLight cytotoxicity assay

T-cell cytotoxicity against UV-killed Mtb-treated target cells was measured using a ToxiLight assay (Lonza), which was carried out as per the manufacturer’s instructions. Briefly, 20μl of media from each well was transferred to a luminometer compatible 96-well plate, 100μl of ToxiLight reagent was then added. Following a 5-minute incubation period, the plate was read using a Glomax 20/20 Luminometer (Glo-Max Discover; Promega).

#### Lysis assay

Tag-it Violet (Biolegend) stained 5x104 THP1-KO or THP1-CD1c cells were left either uninfected or infected with live Mtb (MOI=1) or stimulated with either 1μg/mL LPS (Sigma) or 1μg/mL Pam3CSK4 (InvivoGen) for 48 hours. These cells were then co-cultured with 2.5 × 10³ CD1c-autoreactive T-cells or cultured alone as matched APC-only controls for 48 hours. Cells were collected, stained with fixable live/dead stain, and analysed by flow cytometry. THP1 target cells were identified as Tag-it Violet-positive cells, and the frequency of dead target cells was determined within this gate. Specific lysis was calculated as the percentage of dead THP1 cells in the T-cell co-culture condition minus the percentage of dead THP1 cells in the matched APC-only control. This normalisation was performed independently for each matched condition within each experiment.

### Luminex xMAP assays

T-cells were re-stimulated overnight with irradiated THP1-KO, or THP1-CD1c cells or left unstimulated. Additionally, CD1c-autoreative T-cells, or control irrelevant T-cells, were co-cultured with untreated or UV-killed Mtb (MOI 10) treated THP1-KO and THP1-CD1c cells. Mtb untreated or treated THP1 cells and T-cells were cultured alone as control. At 24 hours after stimulation, supernatant was removed from each T-cell co-culture for multiplex cytokine analysis, cells were pelleted and the supernatant frozen. We followed manufacturer’s protocol to determine concentrations of cytokines (Life Technologies). A Bioplex 200 platform (Bio-Rad) was used to measure concentrations of cytokines in a multiplex panel: GM-CSF, IFNγ, IL-1β, IL-10, IL-12p70, IL-13, IL-17a, IL-2, IL-22, IL-4, IL-5, IL-8, IP-10, MIP-1α, MIP-1β, Oncostatin M, RANTES and TNFα or the cytokine 35-plex human panel (Thermo Fisher Scientific).

### Mtb Lux growth assays

THP1-KO and THP1-CD1c were infected with Mtb lux (MOI=1). After overnight infection, cells were transferred from vented flasks to 50 mL Falcon tubes after detachment with Versene solution (Sigma Aldrich) for 10 min. Cells were then washed with Hanks’ balanced salt solution (HBSS) without Ca/Mg (Thermo Fisher Scientific) and then centrifuged at 380×g for 5 minutes at 4°C to remove extracellular bacteria. THP1-KO and THP1-CD1c were next resuspended in T-cell media containing kanamycin (25 μg/mL), co-cultured with T-cells and incubated at 37°C, 21% O_2_ and 5% CO_2_. Mtb growth was monitored by luminescence (GloMax 20/20 Luminometer, Promega) for 7 days. Luminescence values reflect viable lux-expressing Mtb under the assay conditions and do not distinguish intracellular from extracellular bacteria. Relative luminescence values for each condition were normalised to the matched infected APC-only control within each experimental replicate, allowing comparison of net bacterial burden across co-culture conditions.

### Single cell library preparation and sequencing

CD3-enriched PBMCs were washed in cold staining buffer (1% BSA in PBS), counted, and assessed for viability. One million cells per sample were resuspended in binding buffer containing 50nM dasatinib (Sigma), blocked (as above) and then stained with PE-dCODE™ dextramers conjugated to CD1c-endo, in combination with the following fluorochrome-conjugated antibodies: FITC-conjugated anti-human CD3 (clone UCHT1) and viability dye (Live/Dead Near IR, Invitrogen, Basingstoke, UK). Following a 30-minute incubation at 4 °C, cells were washed twice with ice-cold PBS supplemented with 1% FBS and sorted on a BD FACSAria cell sorter (BD Biosciences). Sorted cell fractions were labelled with sample-specific TotalSeq™-C hashtag antibodies. Flow cytometry data were analysed using FlowJo v10.8.1 (FlowJo LLC). Sorted cells were loaded onto a 10x Genomics Chromium Chip G at a target of ∼45,000 cells per lane using the Single Cell V(D)J Reagent Kit v1.1. First-strand cDNA synthesis was performed according to the manufacturer’s protocol. After emulsion disruption and nucleic acid recovery, cDNA amplification proceeded following the Chromium Next GEM Single Cell V(D)J v1.1 user guide.

### scRNA-seq alignment and analysis of CD1c-endo dextramer-sorted T-cells

Sequencing reads were de-multiplexed and aligned using Cell Ranger v7.2.0, with mapping to the GRCh38 reference genome (build 2020-A, 10x Genomics). A total of 14,524 single CD3⁺ T-cells were initially profiled by scRNA-seq following dextramer-guided cell sorting. Cells were sorted into CD3⁺CD1c-endo-positive and CD3⁺CD1c-endo-negative fractions using dCODE™ dextramer reagents. Raw gene expression matrices were generated and processed for downstream analysis.

### Quality control and filtering

Cells underwent rigorous quality control (QC) to ensure data integrity and remove low-quality or ambiguous profiles. Specifically, the following metrics were calculated per cell: number of detected genes, total unique molecular identifiers, mitochondrial gene content, and ribosomal gene content. Additionally, two independent doublet detection methods were applied: Scrublet, and scDblFinder. Outlier cells were excluded based on the following criteria: low gene content and Unique Molecular Identifier (UMI) counts, high mitochondrial content (>10%) and high ribosomal content (>50%). Predicted doublets from at least one of the two detection tools. After filtering, a total of 11,804 high-quality single T-cells were retained for downstream analysis.

### Annotation of CD1c-endo binding

Cells were annotated as CD1c-endo-positive or -negative based on dextramer binding. Of the 11,804 that passed quality control, 9,518 lacked detectable dextramer binding and were initially labelled as CD1c-endo negative. These were further stratified by sorting origin: 7,652 cells originated from the CD1c-endo-positive sort gate, and 1,866 cells came from the CD1c-endo-negative sort gate. Only the 1,866 cells from the CD1c-endo-negative sort gate were retained as confidently assigned CD1c-endo-negative T-cells. The remaining 7,652 cells, despite lacking dextramer signal, were excluded due to possible false negatives introduced during staining or sorting. To accurately classify CD1c-endo-positive cells, a density-based thresholding approach was applied. The distribution of dextramer signal was modelled using kernel density estimation (KDE). To define confidently positive cells, thresholds were set based on the 1st and 99th percentiles and the interquartile range (IQR) of dextramer intensity. Cells with signal below the lower bound (Q1 – 1.5 × IQR) were excluded as potential noise or non-specific binders. This approach minimised false positives and enabled robust separation of CD1c-endo-positive and -negative populations. After applying quality control filters and dextramer-based gating, CD1c-endo-positive T-cells were defined as those confidently binding dextramer above the KDE threshold, while CD1c-endo-negative T-cells comprised the 1,866 cells from the negative gate lacking any detectable dextramer signal.

### Subsampling analysis to assess robustness

To control for group size imbalance (CD1c-endo⁺ vs CD1c-endo⁻), we performed a subsampling analysis. From each group, a random subset of cells equal to the smallest group size (1,866) was sampled using np.random.choice (without replacement). This balanced dataset was reanalysed to ensure that key differential gene expression patterns and clustering results were consistent across equal-sized groups. The findings from the full dataset were confirmed, indicating that group size imbalance did not bias the results, validating the robustness of the conclusions.

### Study approval

All clinical studies were conducted according to the Declaration of Helsinki principles. All participants gave written informed consent. Paraffin-embedded lung tissue from TB patients or those with adenocarcinoma were retrieved from the histology archive at University Hospital Southampton with approval by the Institutional Review Board (Reference 12/NW/0794: SRB04_14). The ethics committee approved immunohistochemical analysis without individual informed consent since it was surplus archived tissue taken as part of routine care.

#### Data availability

scRNA-seq data have been deposited in the European Nucleotide Archive (ENA) under accession number PRJEB94332 and are publicly available as of the date of publication. Additional data and analysis code are available from the corresponding author upon request.

#### Statistical Analysis

GraphPad Prism version 10 (GraphPad Software, Inc.) was used for statistical analysis, and P values ≤0.05 were considered statistically significant. The Mann–Whitney U test, T tests, one-way ANOVA, Wilcoxon signed-rank test, 2-way ANOVA and Tukeys test were used as stated in the figure legends.

## Acknowledgements

We thank Richard Jewell, Carolann McGuire, and Sarah Pearson for their assistance with flow cytometry (FACS facility, Faculty of Medicine, University of Southampton), Jon Ward (Histochemistry Research Unit) for undertaking the immunohistochemical staining. We also thank Regina Teo for her support in managing our departmental laboratory section. We thank Lauren Chessum and Shaliny Ramachandran at Immunocore for assistance with single-cell TCR sequencing. M.M and A.L were supported by a UKRI MRC DTP studentship award (MR/W007045/1). S.H.F was supported by a MRC iCase studentship and Immunocore Ltd. This work was supported by a Public Health England funded PhD studentship and a University of Southampton Vice Chancellor award to J.G and K.N. A.L was supported by the Wellcome Trust (210662/Z/18/Z) and the Bill and Melinda Gates Foundation (OPP1137006). D.B was supported by a studentship funded by the Institute for Life Sciences, University of Southampton. PE was supported by MRC (MR/P023754/1 and MR/W025728/1). S.M was supported by MRC (MR/S024220/1) and Cancer Research UK (23562). The UK funded award (MR/S024220/1) is part of the EDCTP2 programme supported by the European Union. We acknowledge the support of the Southampton National Institute for Health Research Biomedical Research Centre.

## Author contributions

Joint first author order reflects contribution to data generation. M.M., S.H.F., D.G.B and S.M. designed research; M.M., S.H.F., J.G., D.G.B., K.N., D.B., S.S., R.S-K., P.T-S., A.L., R.S., A.V., L.T., A.W., L.D., P.E., and S.M. performed research; R.S-K., A.L., M.L., D.K.C., S.S., A.L., A.V., and P.E. contributed new reagents/analytic tools; M.M., S.H.F., J.G., D.G.B., L.T., A.W., M.L., L.D., D.K.C., A.L., S.S., L.T., A.L., A.V., P.E., and S.M. analyzed data; and S.M. wrote the paper, all authors edited and approved final version.

## Supplementary Figures

**Figure S1.**
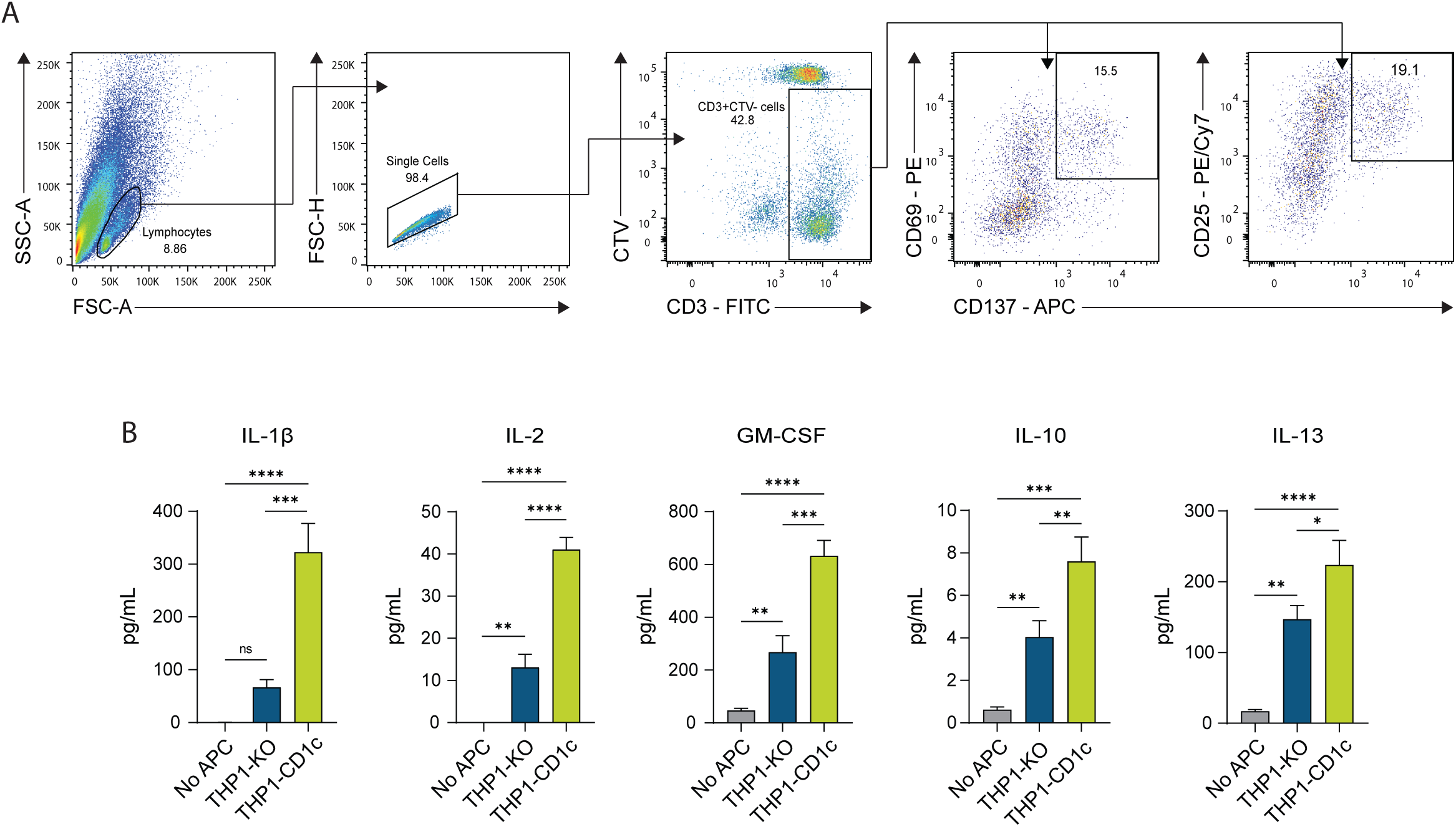
(A) Flow cytometry gating strategy analysing the expression of the T-cell activation markers CD69, CD25, and CD137 on proliferated CD3^+^CTV^-^ T-cells. (B) Cytokine release by T-cells first expanded with THP1-CD1c APCs and then stimulated overnight with THP1-KO or THP1-CD1c APCs. Cytokine secretion was measured by Luminex array. * P < 0.05; ** P < 0.01, *** P < 0.001, **** P < 0.0001 (one-way ANOVA with Tukey’s multiple comparison test). Mean and SD of triplicate measurements are shown and are representative of three individual donors.

**Figure S2.**
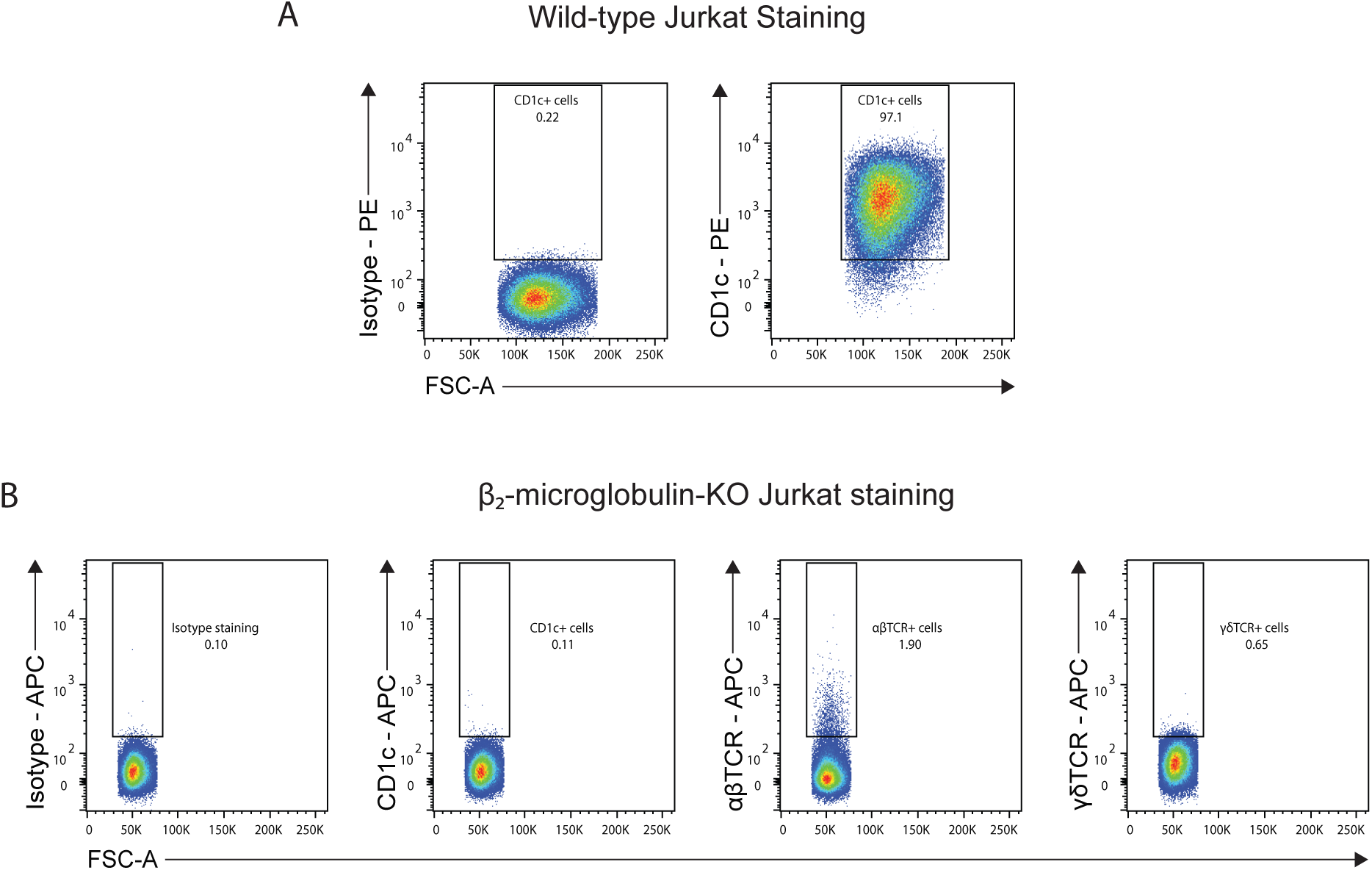
(A) Flow cytometry gating strategy of live HLA-DR^+^/CD11c^+^ MoDCs (top), and line graphs showing the effect of Mtb infection on the expression of CD1a, CD1b, and CD1c on MoDCs (bottom). (B) Flow cytometry gating strategy depicting live CD14^+^ THP1-CD1c cells stained with anti-CD1c antibody.

**Figure S3.**
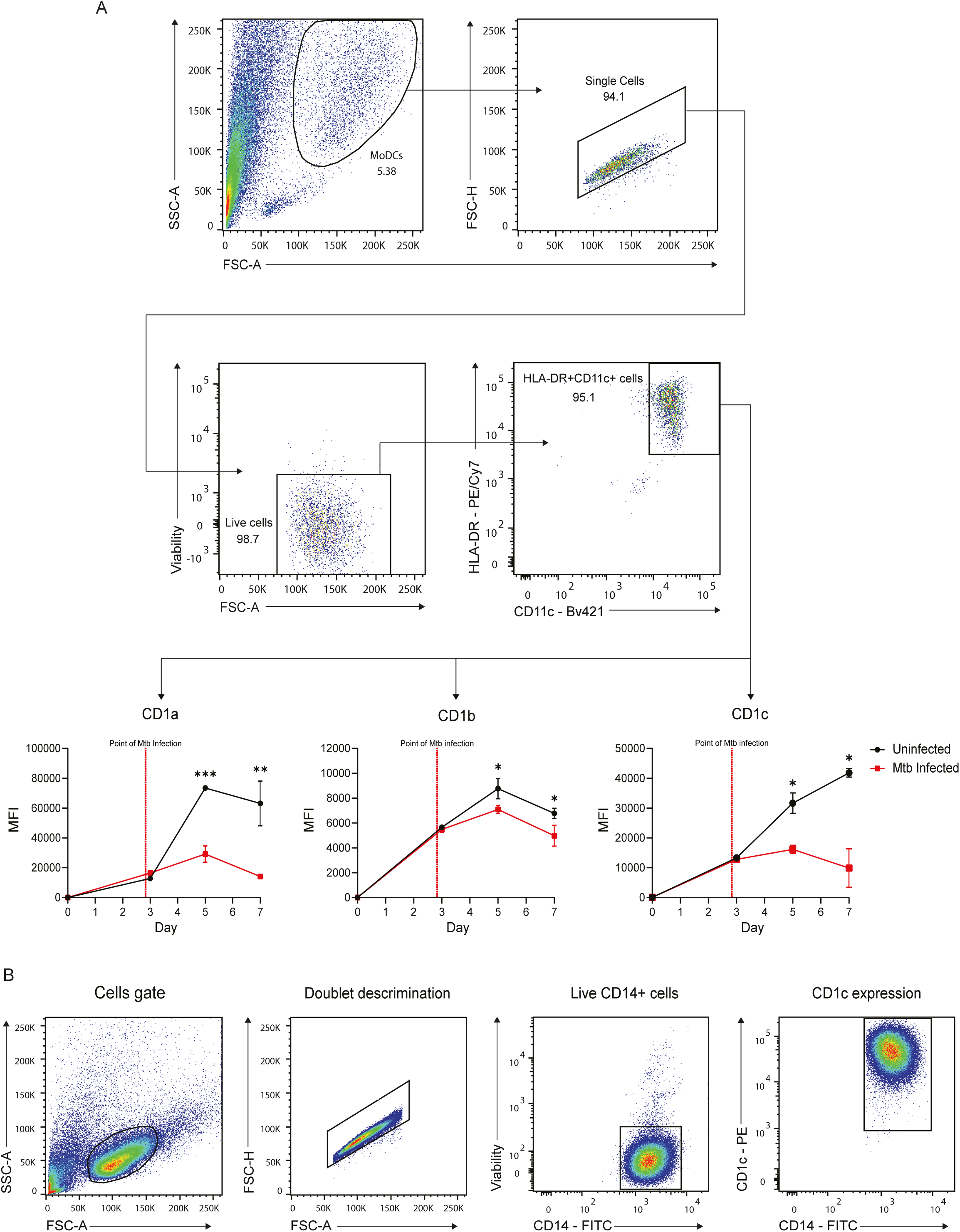
Flow cytometry gating strategy depicting live CD3^+^ T-cells comprising the negative (cells that did not bind the streptamers) or the CD1c-endo streptamer positive T-cell fraction (containing 1.14% CD1c-endo positive T-cells) stained with CD1c-endo dextramers. CD1c-endo dextramer positive T-cells were sorted and expanded. After expansion, T-cells were either unstained, or stained with an irrelevant tetramer or with CD1c-endo tetramer. Enriched T-cells brightly stain with CD1c-endo tetramers.

**Figure S4.**
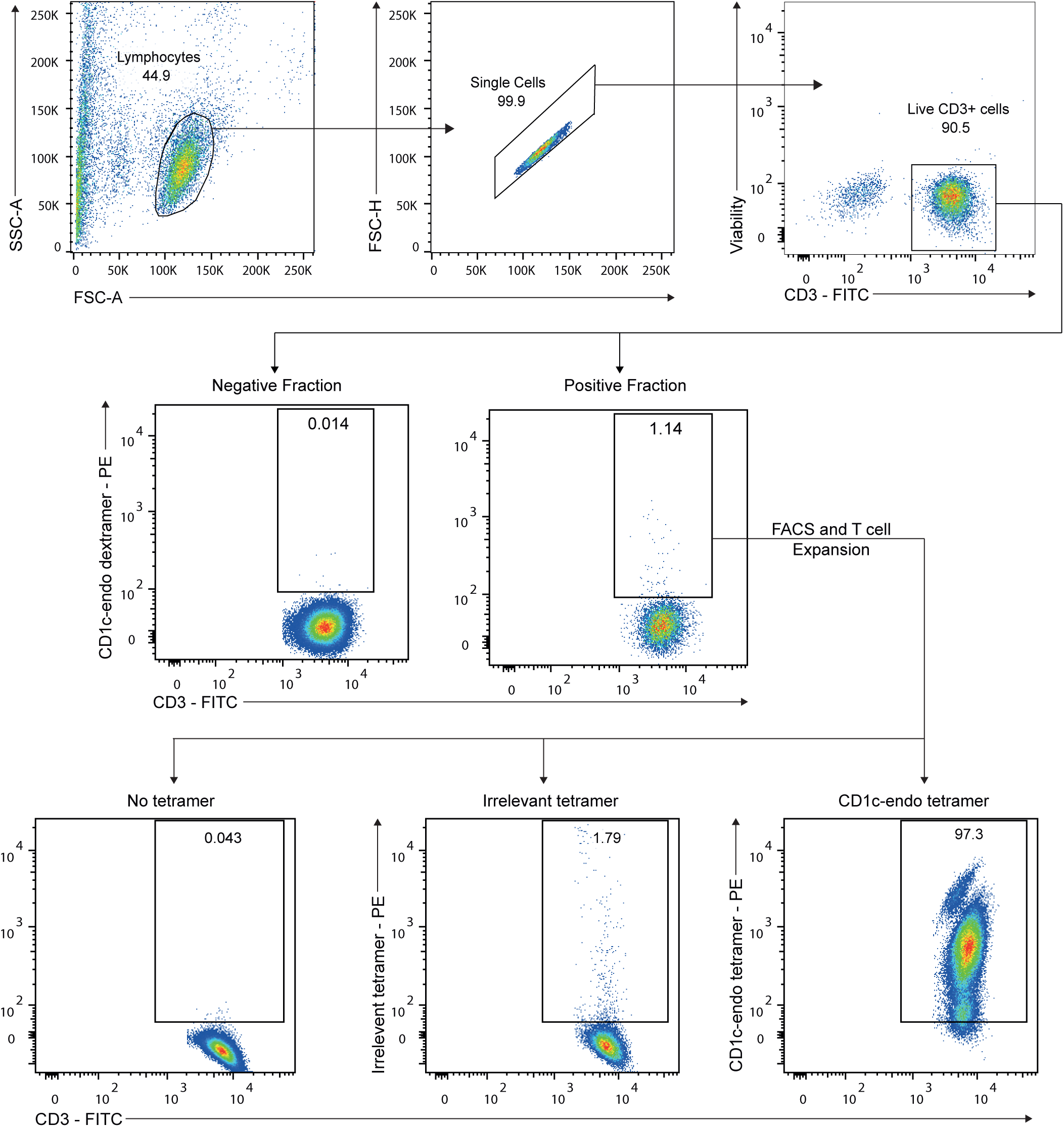
(A) Flow cytometry dot plots showing high expression of CD1c on wild-type JRT3.5 Jurkat T-cells. (B) Flow cytometry dot plots showing the absence of CD1c and TCR expression on β2-microglobulin knock-out JRT3.5 Jurkat T-cells.

**Figure S5.**
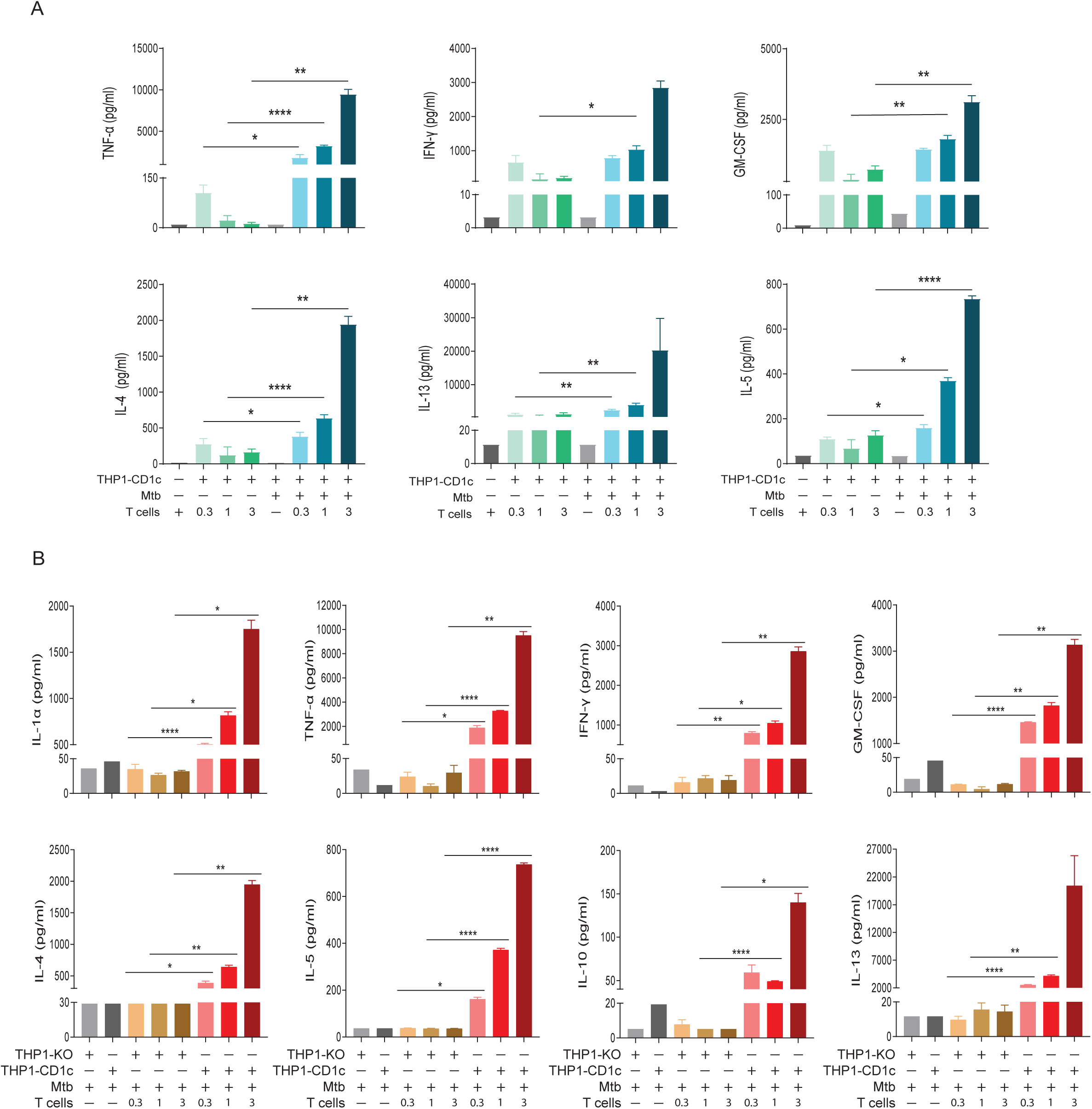
(A) CD1c-autoreactive T-cells secrete cytokines in response to untreated THP1-CD1c APCs in an autoreactive manner. Importantly, CD1c-autoreactive T-cells release significantly higher concentrations of cytokine when cultured with Mtb-infected THP1-CD1c APCs. (B) CD1c-autoreactive T-cells secrete diverse cytokines in a CD1c dependent manner. CD1c-autoreactive T-cells release cytokines when cultured with Mtb-treated THP1-CD1c APCs but not when were cultured with Mtb-treated THP1-KO APCs. Data are representative of two independent experiments, each preformed in triplicate. * P < 0.05; ** P < 0.01, **** P < 0.0001 (Two-way ANOVA).

**Figure S6.**
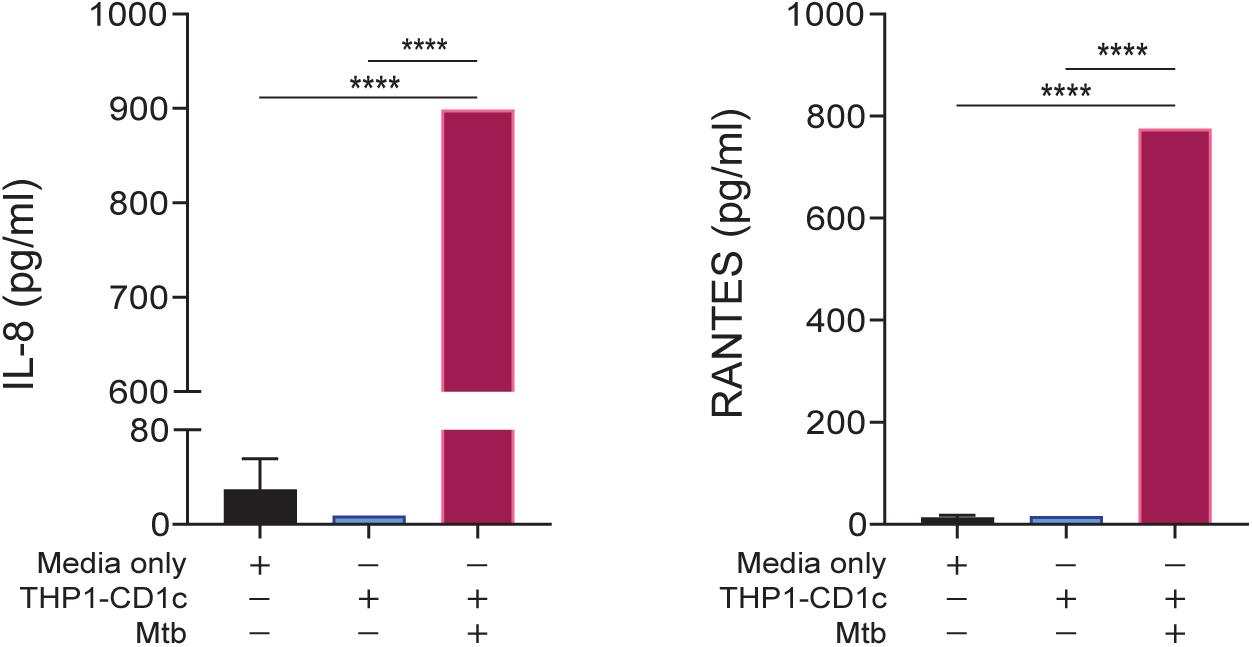
UV Mtb-treated THP1-CD1c APCs secrete chemokines IL-8 and RANTES (pink bars), in comparison to media only (black bars), and untreated THP1-CD1c APCs (blue bars). Data are representative of two independent experiments, each preformed in triplicate. **** P < 0.0001 (Two-way ANOVA).

**Figure S7.**
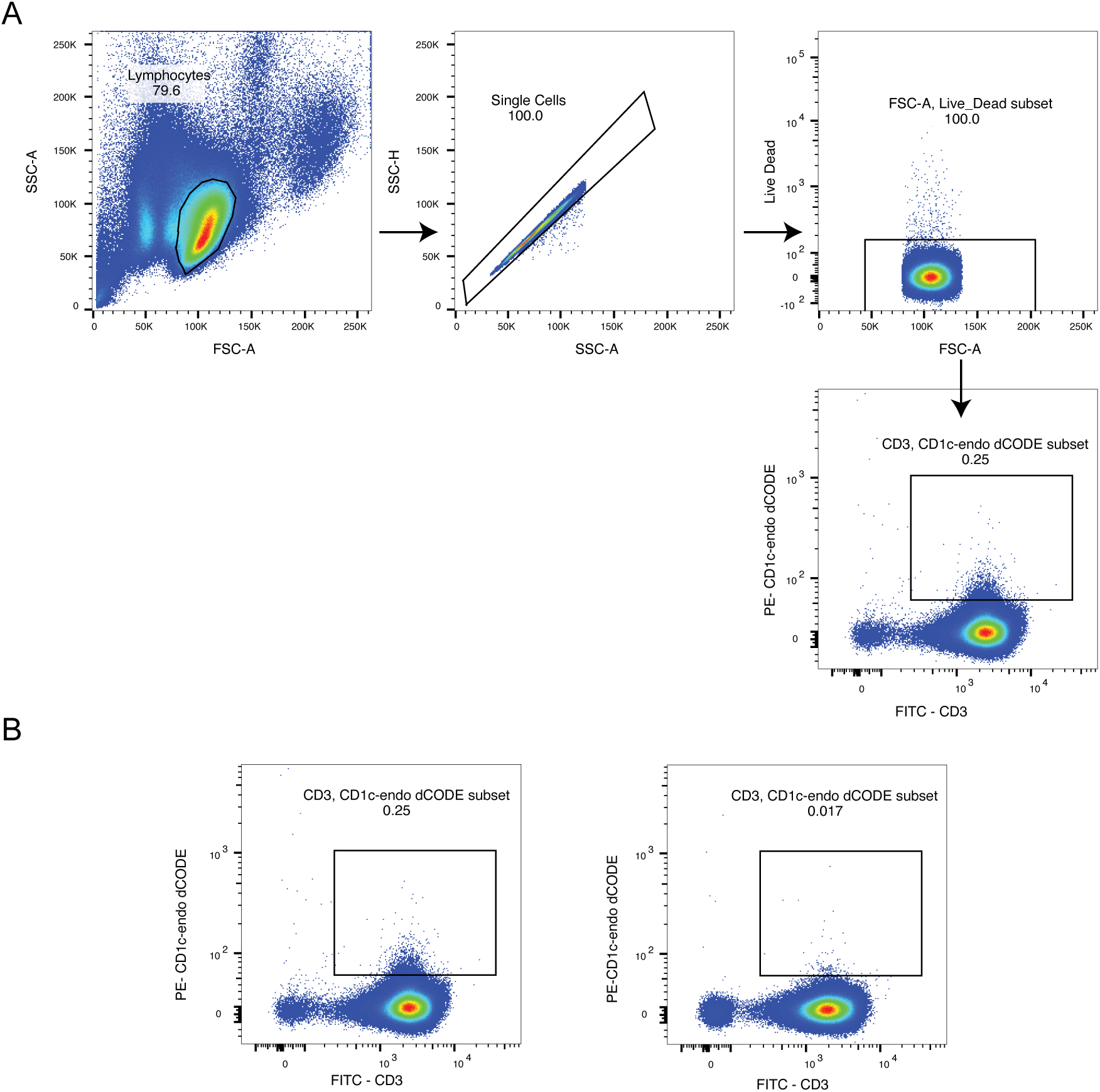
(A) Representative flow cytometry gating strategy used to isolate CD3⁺ T-cells stained with CD1c-endo dCODE dextramers. (B) Representative plots from two donors showing distinct CD3⁺CD1c-endo⁺ and CD3⁺CD1c-endo⁻ T-cell populations. Percentages indicate the proportion of positive cells within the CD3⁺ gate.

**Figure S8.**
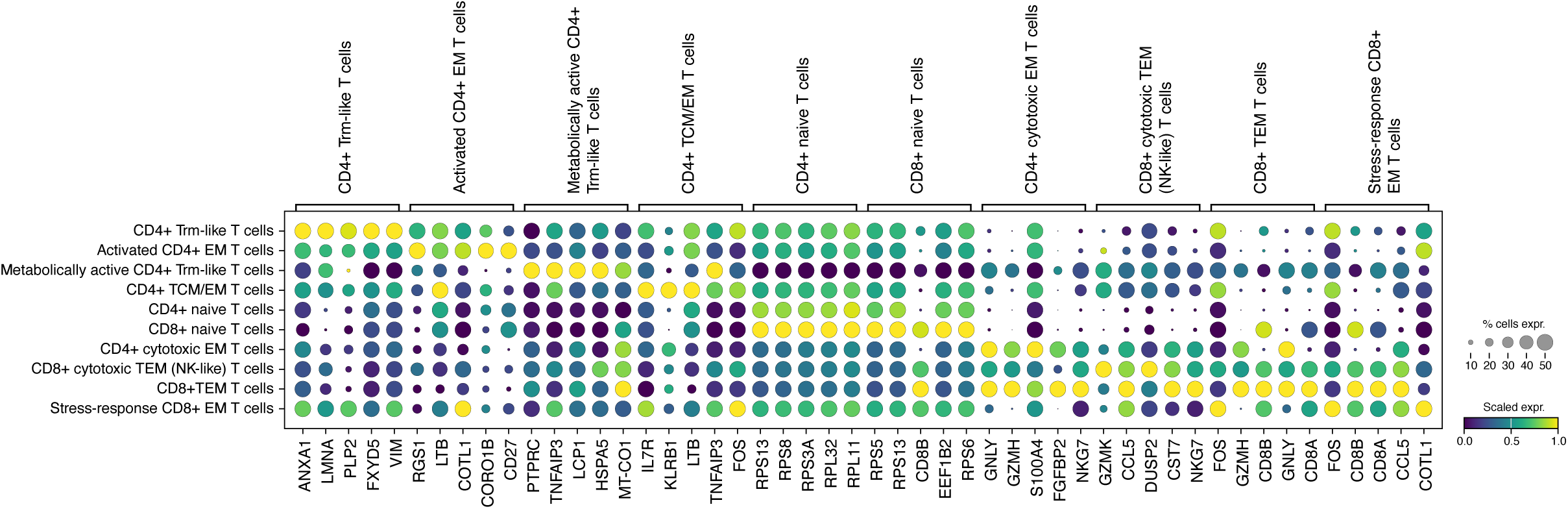
Dot plot showing the top 5 ranked marker genes for each annotated T-cell population, derived from all high-quality T-cells (n = 11,804). Dot size represents the proportion of cells expressing each gene within the corresponding cluster, while colour indicates the scaled average expression (z-score). This visualization highlights the transcriptional signatures that define distinct T-cell subsets, including naïve, memory, effector, and metabolically active populations.

**Figure S9.**
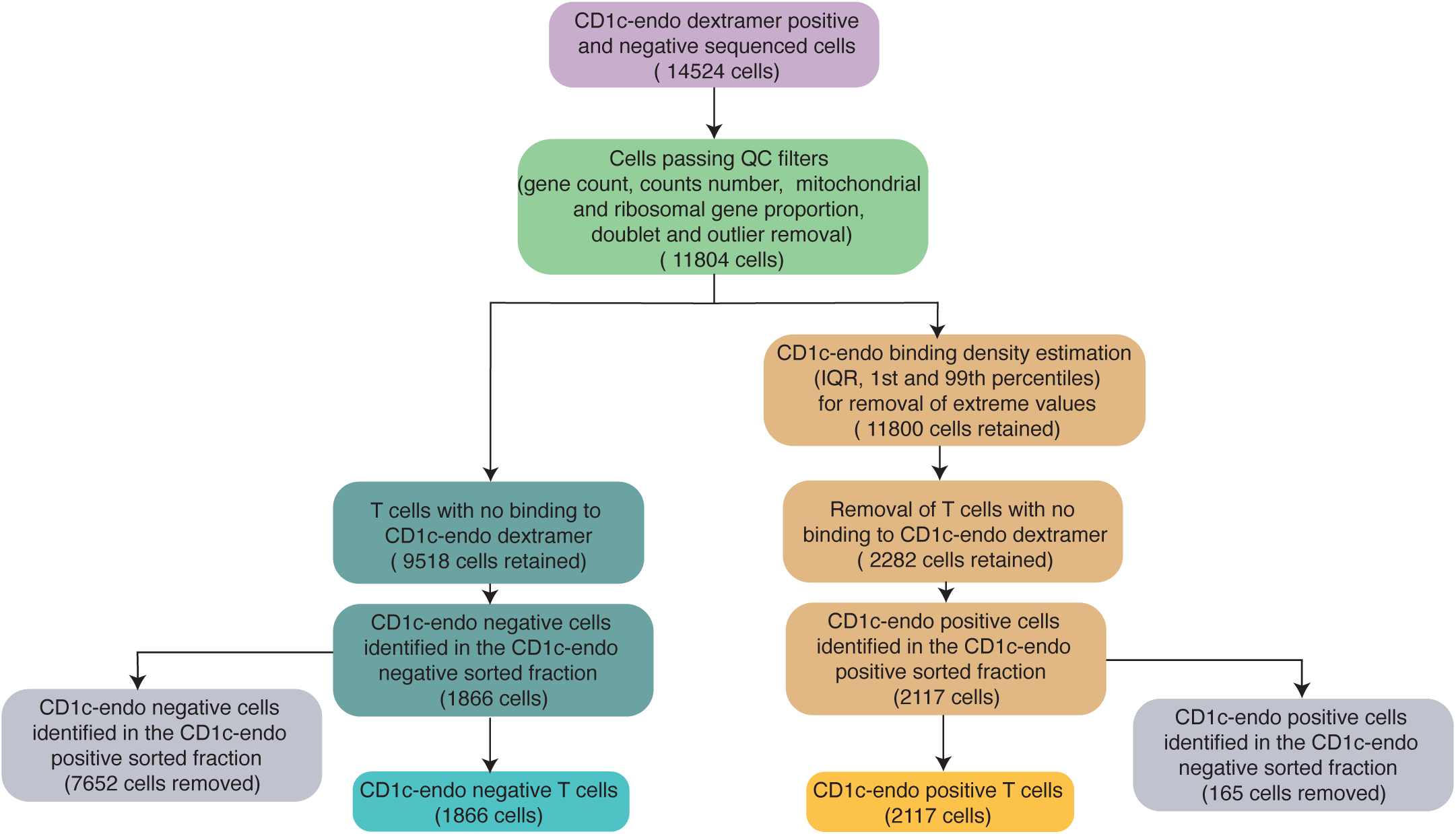
Filtering strategy for identifying CD1c-endo dextramer-bound T-cells in scRNA-seq data from sorted populations: A total of 14,524 cells were initially profiled by scRNA-seq from dextramer-sorted fractions. Following standard quality control, including thresholds for gene and UMI counts, mitochondrial and ribosomal content, doublet detection, and outlier removal, 11,804 high-quality cells were retained. CD1c-endo-negative cells (n = 9,518) were defined by the absence of detectable dextramer signal and further stratified by their sort gate: 7,652 originated from the CD1c-endo-positive gate and 1,866 from the CD1c-endo-negative gate. To minimise false negatives, only the 1,866 cells from the negative sort gate were retained as confidently CD1c-endo-negative. To define CD1c-endo-positive cells, dextramer signal intensities were modelled using kernel density estimation (KDE) and quantile-based filtering (1st and 99th percentiles, IQR). Cells with signal below the lower bound (Q1 – 1.5×IQR) or extreme values were excluded, yielding 11,800 cells. Of these, 2,288 cells with no binding signal were removed. Among the remaining 2,282 CD1c-endo-positive candidates, 2,117 were from the CD1c-endo-positive sort gate and 165 from the negative gate. To ensure specificity, only the 2,117 cells from the positive sort gate were retained as confidently CD1c-endo-positive. This combined quality control and gating strategy enabled robust identification of CD1c-endo-binding T-cells in the single-cell dataset.

**Table S1.**
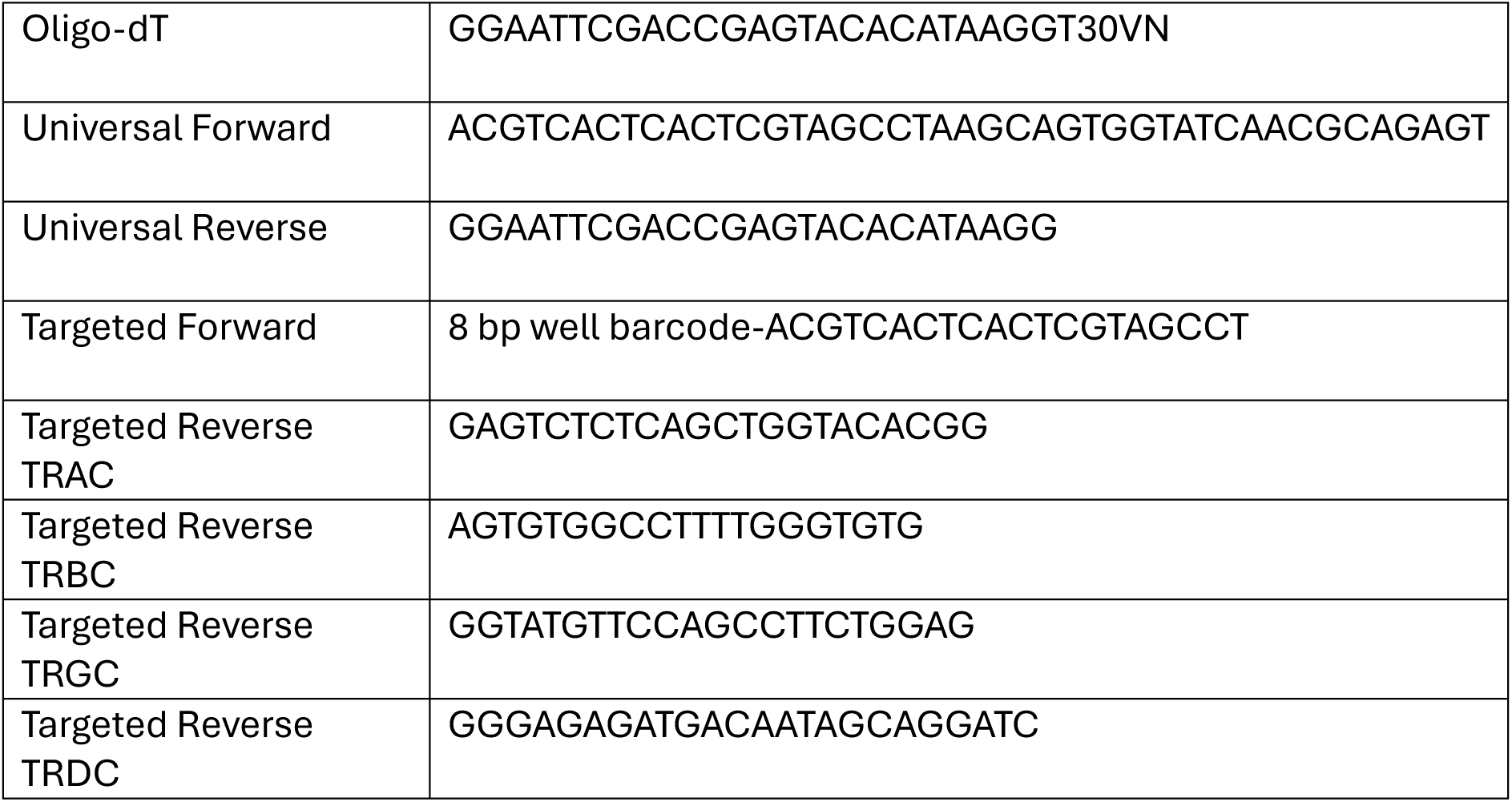
Primer sequences used for targeted single-cell TCR sequencin. Sequences are shown for the oligo-dT primer, universal forward and reverse primers, targeted forward primer containing the 8-bp well barcode, and reverse primers specific for TRAC, TRBC, TRGC and TRDC constant regions.

## References

1 Wallis, R. S. et al. Tuberculosis--advances in development of new drugs, treatment regimens, host-directed therapies, and biomarkers. Lancet Infect Dis 16, e34–46 (2016). 10.1016/S1473-3099(16)00070-0

2 Programme, G. T. Global Tuberculosis Report, 2021).

3 Hui, D. et al. Global tuberculosis response off track: urgent priorities to end the world’s top infectious killer. Lancet (2025). 10.1016/S0140-6736(25)02433-X

4 Nguipdop-Djomo, P., Heldal, E., Rodrigues, L. C., Abubakar, I. & Mangtani, P. Duration of BCG protection against tuberculosis and change in effectiveness with time since vaccination in Norway: a retrospective population-based cohort study. Lancet Infect Dis 16, 219–226 (2016). 10.1016/S1473-3099(15)00400-4

5 O’Garra, A. et al. The immune response in tuberculosis. Annu Rev Immunol 31, 475–527 (2013). 10.1146/annurev-immunol-032712-095939

6 Jasenosky, L. D., Scriba, T. J., Hanekom, W. A. & Goldfeld, A. E. T cells and adaptive immunity to Mycobacterium tuberculosis in humans. Immunol Rev 264, 74–87 (2015). 10.1111/imr.12274

7 Ndiaye, B. P. et al. Safety, immunogenicity, and efficacy of the candidate tuberculosis vaccine MVA85A in healthy adults infected with HIV-1: a randomised, placebo-controlled, phase 2 trial. The lancet. Respiratory medicine 3, 190-200 (2015). 10.1016/S2213-2600(15)00037-5

8 Tameris, M. D. et al. Safety and efficacy of MVA85A, a new tuberculosis vaccine, in infants previously vaccinated with BCG: a randomised, placebo-controlled phase 2b trial. Lancet 381, 1021–1028 (2013). 10.1016/S0140-6736(13)60177-4

9 Schrager, L. K., Vekemens, J., Drager, N., Lewinsohn, D. M. & Olesen, O. F. The status of tuberculosis vaccine development. Lancet Infect Dis (2020). 10.1016/S1473-3099(19)30625-5

10 Tait, D. R. et al. Final Analysis of a Trial of M72/AS01E Vaccine to Prevent Tuberculosis. N Engl J Med 381, 2429–2439 (2019). 10.1056/NEJMoa1909953

11 Joosten, S. A. et al. Harnessing donor unrestricted T-cells for new vaccines against tuberculosis. Vaccine (2019). 10.1016/j.vaccine.2019.04.050

12 Nunes-Alves, C. et al. In search of a new paradigm for protective immunity to TB. Nat Rev Microbiol 12, 289–299 (2014). 10.1038/nrmicro3230

13 Ruibal, P., Voogd, L., Joosten, S. A. & Ottenhoff, T. H. M. The role of donor-unrestricted T-cells, innate lymphoid cells, and NK cells in anti-mycobacterial immunity. Immunol Rev 301, 30–47 (2021). 10.1111/imr.12948

14 Chancellor, A., Gadola, S. D. & Mansour, S. The versatility of the CD1 lipid antigen presentation pathway. Immunology 154, 196–203 (2018). 10.1111/imm.12912

15 Mori, L., Lepore, M. & De Libero, G. The Immunology of CD1- and MR1-Restricted T Cells. Annu Rev Immunol 34, 479–510 (2016). 10.1146/annurev-immunol-032414-112008

16 Melian, A., Geng, Y. J., Sukhova, G. K., Libby, P. & Porcelli, S. A. CD1 expression in human atherosclerosis. A potential mechanism for T cell activation by foam cells. Am J Pathol 155, 775–786 (1999). 10.1016/S0002-9440(10)65176-0

17 de Jong, A. et al. CD1c presentation of synthetic glycolipid antigens with foreign alkyl branching motifs. Chem Biol 14, 1232–1242 (2007). 10.1016/j.chembiol.2007.09.010

18 Moody, D. B. et al. CD1c-mediated T-cell recognition of isoprenoid glycolipids in Mycobacterium tuberculosis infection. Nature 404, 884–888 (2000). 10.1038/35009119

19 Matsunaga, I. et al. Mycobacterium tuberculosis pks12 produces a novel polyketide presented by CD1c to T cells. J Exp Med 200, 1559–1569 (2004). 10.1084/jem.20041429

20 Scharf, L. et al. The 2.5 A structure of CD1c in complex with a mycobacterial lipid reveals an open groove ideally suited for diverse antigen presentation. Immunity 33, 853–862 (2010). 10.1016/j.immuni.2010.11.026

21 Roy, S. et al. Molecular basis of mycobacterial lipid antigen presentation by CD1c and its recognition by alphabeta T cells. Proc Natl Acad Sci U S A 111, E4648–4657 (2014). 10.1073/pnas.1408549111

22 Ly, D. et al. CD1c tetramers detect ex vivo T cell responses to processed phosphomycoketide antigens. J Exp Med 210, 729–741 (2013). 10.1084/jem.20120624

23 de Lalla, C. et al. High-frequency and adaptive-like dynamics of human CD1 self-reactive T cells. Eur J Immunol 41, 602–610 (2011). 10.1002/eji.201041211

24 de Jong, A. et al. CD1a-autoreactive T cells are a normal component of the human alphabeta T cell repertoire. Nat Immunol 11, 1102–1109 (2010). 10.1038/ni.1956

25 Wun, K. S. et al. T cell autoreactivity directed toward CD1c itself rather than toward carried self lipids. Nat Immunol 19, 397–406 (2018). 10.1038/s41590-018-0065-7

26 Mansour, S. et al. Cholesteryl esters stabilize human CD1c conformations for recognition by self-reactive T cells. Proc Natl Acad Sci U S A 113, E1266–1275 (2016). 10.1073/pnas.1519246113

27 Lepore, M. et al. A novel self-lipid antigen targets human T cells against CD1c(+) leukemias. J Exp Med 211, 1363–1377 (2014). 10.1084/jem.20140410

28 Sieling, P. A. et al. Human double-negative T cells in systemic lupus erythematosus provide help for IgG and are restricted by CD1c. J Immunol 165, 5338–5344 (2000). 10.4049/jimmunol.165.9.5338

29 Roura-Mir, C. et al. CD1a and CD1c activate intrathyroidal T cells during Graves’ disease and Hashimoto’s thyroiditis. J Immunol 174, 3773–3780 (2005). 10.4049/jimmunol.174.6.3773

30 Suzuki, T. A. et al. Codiversification of gut microbiota with humans. Science 377, 1328–1332 (2022). 10.1126/science.abm7759

31 Brites, D. & Gagneux, S. Co-evolution of Mycobacterium tuberculosis and Homo sapiens. Immunol Rev 264, 6–24 (2015). 10.1111/imr.12264

32 Felio, K. et al. CD1-restricted adaptive immune responses to Mycobacteria in human group 1 CD1 transgenic mice. J Exp Med 206, 2497–2509 (2009). 10.1084/jem.20090898

33 Li, S., Choi, H. J., Felio, K. & Wang, C. R. Autoreactive CD1b-restricted T cells: a new innate-like T-cell population that contributes to immunity against infection. Blood 118, 3870–3878 (2011). 10.1182/blood-2011-03-341941

34 Van Rhijn, I. et al. Human autoreactive T cells recognize CD1b and phospholipids. Proc Natl Acad Sci U S A 113, 380–385 (2016). 10.1073/pnas.1520947112

35 Cotton, R. N., Shahine, A., Rossjohn, J. & Moody, D. B. Lipids hide or step aside for CD1-autoreactive T cell receptors. Curr Opin Immunol 52, 93–99 (2018). 10.1016/j.coi.2018.04.013

36 Stenger, S., Niazi, K. R. & Modlin, R. L. Down-regulation of CD1 on antigen-presenting cells by infection with Mycobacterium tuberculosis. J Immunol 161, 3582–3588 (1998).

37 Wen, Q. et al. MiR-381-3p Regulates the Antigen-Presenting Capability of Dendritic Cells and Represses Antituberculosis Cellular Immune Responses by Targeting CD1c. J Immunol 197, 580–589 (2016). 10.4049/jimmunol.1500481

38 Pacis, A. et al. Gene activation precedes DNA demethylation in response to infection in human dendritic cells. Proc Natl Acad Sci U S A 116, 6938–6943 (2019). 10.1073/pnas.1814700116

39 Spada, F. M. et al. Self-recognition of CD1 by gamma/delta T cells: implications for innate immunity. J Exp Med 191, 937–948 (2000). 10.1084/jem.191.6.937

40 Brennan, P. J. et al. Invariant natural killer T cells recognize lipid self antigen induced by microbial danger signals. Nat Immunol 12, 1202–1211 (2011). 10.1038/ni.2143

41 Reljic, R., Stylianou, E., Balu, S. & Ma, J. K. Cytokine interactions that determine the outcome of Mycobacterial infection of macrophages. Cytokine 51, 42–46 (2010). 10.1016/j.cyto.2010.04.005

42 Bielecka, M. K. et al. A Bioengineered Three-Dimensional Cell Culture Platform Integrated with Microfluidics To Address Antimicrobial Resistance in Tuberculosis. mBio 8 (2017). 10.1128/mBio.02073-16

43 Karlsson, E. K., Kwiatkowski, D. P. & Sabeti, P. C. Natural selection and infectious disease in human populations. Nat Rev Genet 15, 379–393 (2014). 10.1038/nrg3734

44 Fumagalli, M. et al. Signatures of environmental genetic adaptation pinpoint pathogens as the main selective pressure through human evolution. PLoS Genet 7, e1002355 (2011). 10.1371/journal.pgen.1002355

45 Stenger, S. et al. An antimicrobial activity of cytolytic T cells mediated by granulysin. Science 282, 121–125 (1998). 10.1126/science.282.5386.121

46 Gherardin, N. A. et al. CD36 family members are TCR-independent ligands for CD1 antigen-presenting molecules. Sci Immunol 6 (2021). 10.1126/sciimmunol.abg4176

47 Guo, T. et al. A Subset of Human Autoreactive CD1c-Restricted T Cells Preferentially Expresses TRBV4-1(+) TCRs. J Immunol 200, 500–511 (2018). 10.4049/jimmunol.1700677

48 Grailer, J. et al. A Novel Cell-based Luciferase Reporter Platform for the Development and Characterization of T-Cell Redirecting Therapies and Vaccine Development. J Immunother 46, 96–106 (2023). 10.1097/CJI.0000000000000453

49 Vincent, M. S., Xiong, X., Grant, E. P., Peng, W. & Brenner, M. B. CD1a-, b-, and c-restricted TCRs recognize both self and foreign antigens. J Immunol 175, 6344–6351 (2005). 10.4049/jimmunol.175.10.6344

50 Roy, S. et al. Molecular Analysis of Lipid-Reactive Vdelta1 gammadelta T Cells Identified by CD1c Tetramers. J Immunol 196, 1933–1942 (2016). 10.4049/jimmunol.1502202

51 Reinink, P. et al. CD1b presents self and Borrelia burgdorferi diacylglycerols to human T cells. Eur J Immunol 49, 737–746 (2019). 10.1002/eji.201847949

52 Huang, S. et al. CD1 lipidomes reveal lipid-binding motifs and size-based antigen-display mechanisms. Cell (2023). 10.1016/j.cell.2023.08.022

53 Zhao, J. et al. Mycolic acid-specific T cells protect against Mycobacterium tuberculosis infection in a humanized transgenic mouse model. Elife 4 (2015). 10.7554/eLife.08525

54 Aquino, A. et al. Exogenous control of the expression of Group I CD1 molecules competent for presentation of microbial nonpeptide antigens to human T lymphocytes. Clin Dev Immunol 2011, 790460 (2011). 10.1155/2011/790460

55 Gagliardi, M. C. et al. Bacillus Calmette-Guerin shares with virulent Mycobacterium tuberculosis the capacity to subvert monocyte differentiation into dendritic cell: implication for its efficacy as a vaccine preventing tuberculosis. Vaccine 22, 3848–3857 (2004). 10.1016/j.vaccine.2004.07.009

56 Comas, I. et al. Out-of-Africa migration and Neolithic coexpansion of Mycobacterium tuberculosis with modern humans. Nat Genet 45, 1176–1182 (2013). 10.1038/ng.2744

57 Paulson, T. Epidemiology: A mortal foe. Nature 502, S2–3 (2013). 10.1038/502S2a

58 Reichmann, M. T. et al. A co-evolutionary perspective on humans and Mycobacterium tuberculosis in the era of systems biology. Elife 14 (2026). 10.7554/eLife.108175

59 Picelli, S. et al. Full-length RNA-seq from single cells using Smart-seq2. Nat Protoc 9, 171–181 (2014). 10.1038/nprot.2014.006

