## Supplementary material for "Human CD1c-autoreactive T-cells recognise *Mycobacterium tuberculosis*–infected antigen-presenting cells and display cytotoxic effector programmes": Suplementary files

### Figure S1

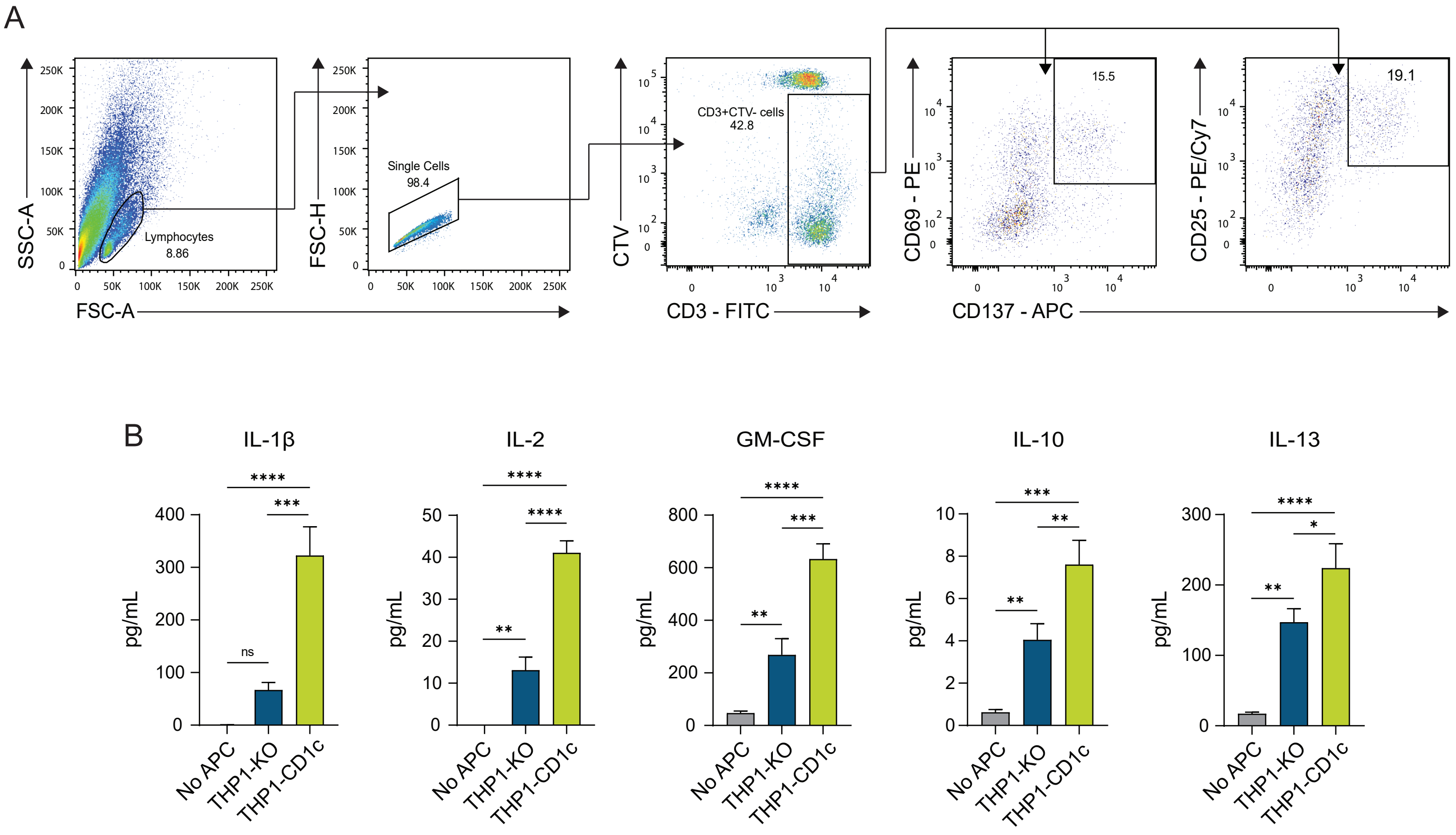


**Fig.S1.** (**A**) Flow cytometry gating strategy analysing the expression of the T-cell activation markers CD69, CD25, and CD137 on proliferated CD3+CTV- T-cells. (**B**) cytokine release by T-cells first expanded with THP1-CD1c APCs and then stimulated overnight with THP1-KO or THP1-CD1c APCs. Cytokine secretion was measured by Luminex array. * $P < 0.05$; ** $P < 0.01$, *** $P < 0.001$, **** $P < 0.0001$ (one-way ANOVA with Tukey's multiple comparison test). Mean and SD of triplicate measurements are shown and are representative of three individual donors.

### Figure S2

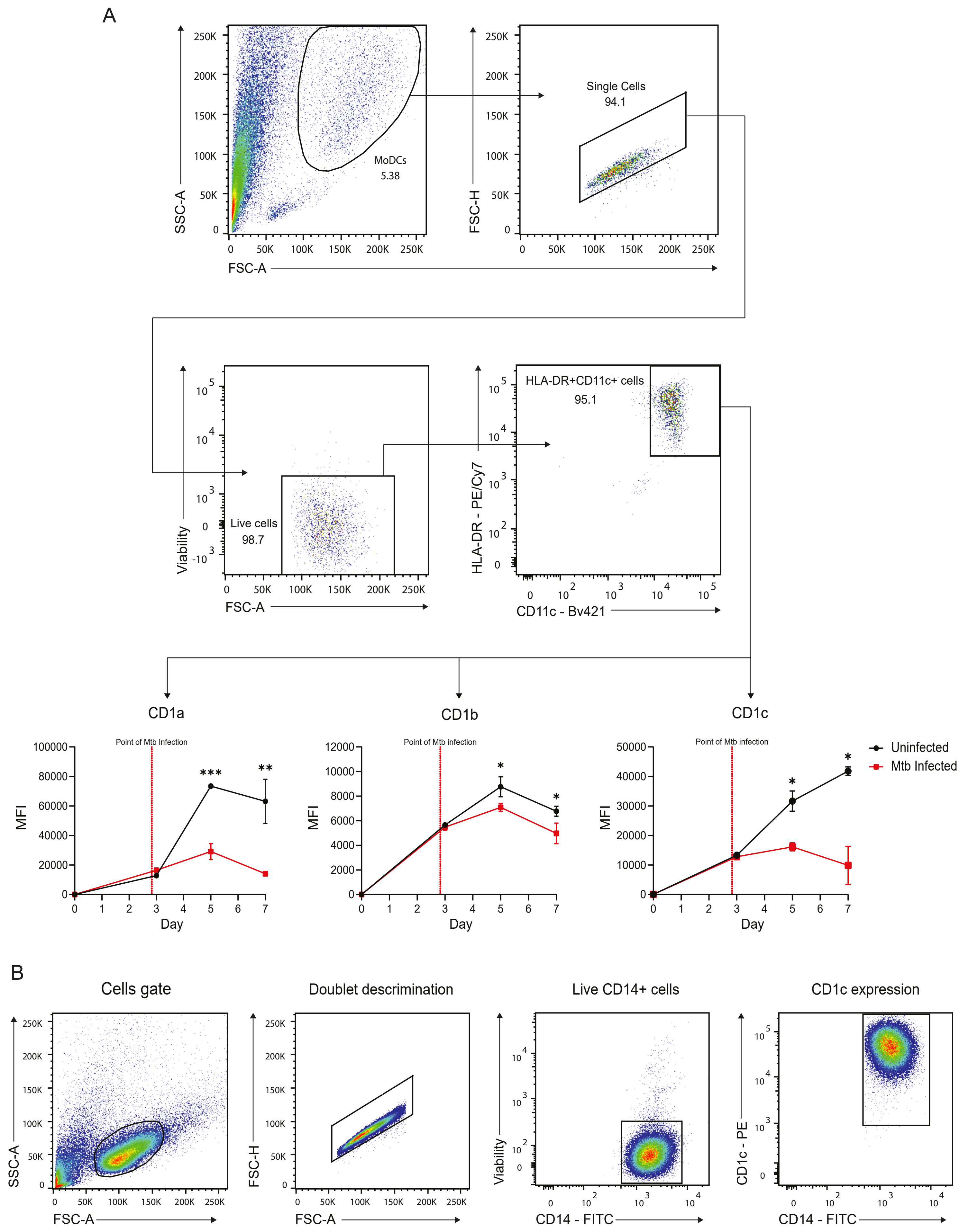
A
MoDCs
5.38
SSC-A
FSC-A
Single Cells
94.1
FSC-H
Live cells
98.7
Viability
HLA-DR+CD11c+ cells
95.1
HLA-DR - PE/Cy7
CD11c - Bv421
CD1a
CD1b
CD1c
Point of Mtb Infection
Point of Mtb infection
MFI
Day
Uninfected
Mtb Infected
B
Cells gate
Doublet descrimination
Live CD14+ cells
CD1c expression
CD14 - FITC
CD1c - PE

**Fig.S2.** (**A**) Flow cytometry gating strategy of live HLA-DR+/CD11c+ MoDCs (top), and line graphs showing the effect of Mtb infection on the expression of CD1a, CD1b, and CD1c on MoDCs (bottom). (**B**) Flow cytometry gating strategy depicting live CD14+ THP1-CD1c cells stained with anti-CD1c antibody.

### Figure S3

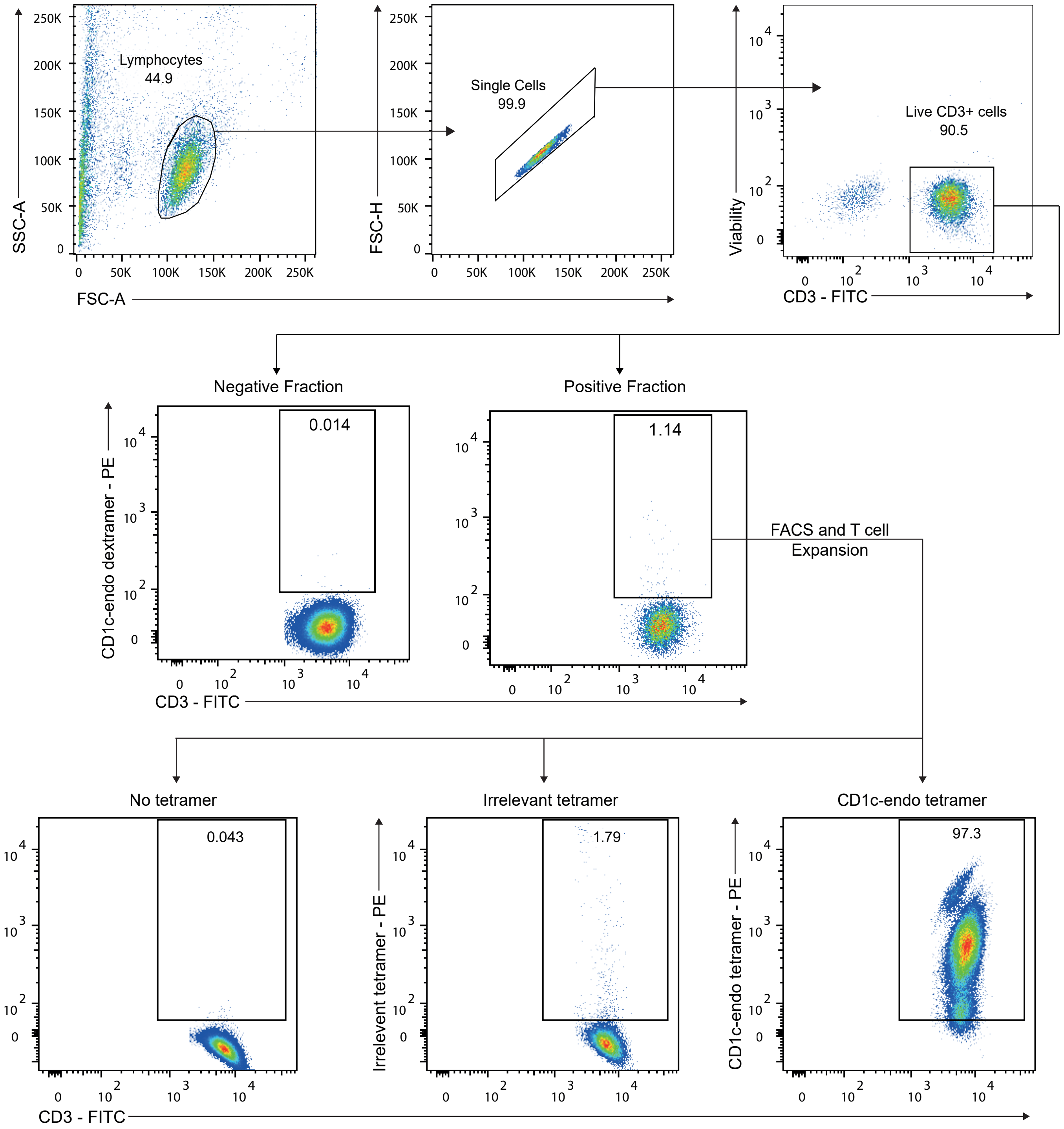


**Fig.S3.** Flow cytometry gating strategy depicting live CD3+ T-cells comprising the negative (cells that did not bind the streptamers) or the CD1c-endo streptamer positive T-cell fraction (containing 1.14% CD1c-endo positive T-cells) stained with CD1c-endo dextramers. CD1c-endo dextramer positive T-cells were sorted and expanded. After expansion, T-cells were either unstained, or stained with an irrelevant tetramer or with CD1c-endo tetramer. Enriched T-cells brightly stain with CD1c-endo tetramers.

### Figure S4

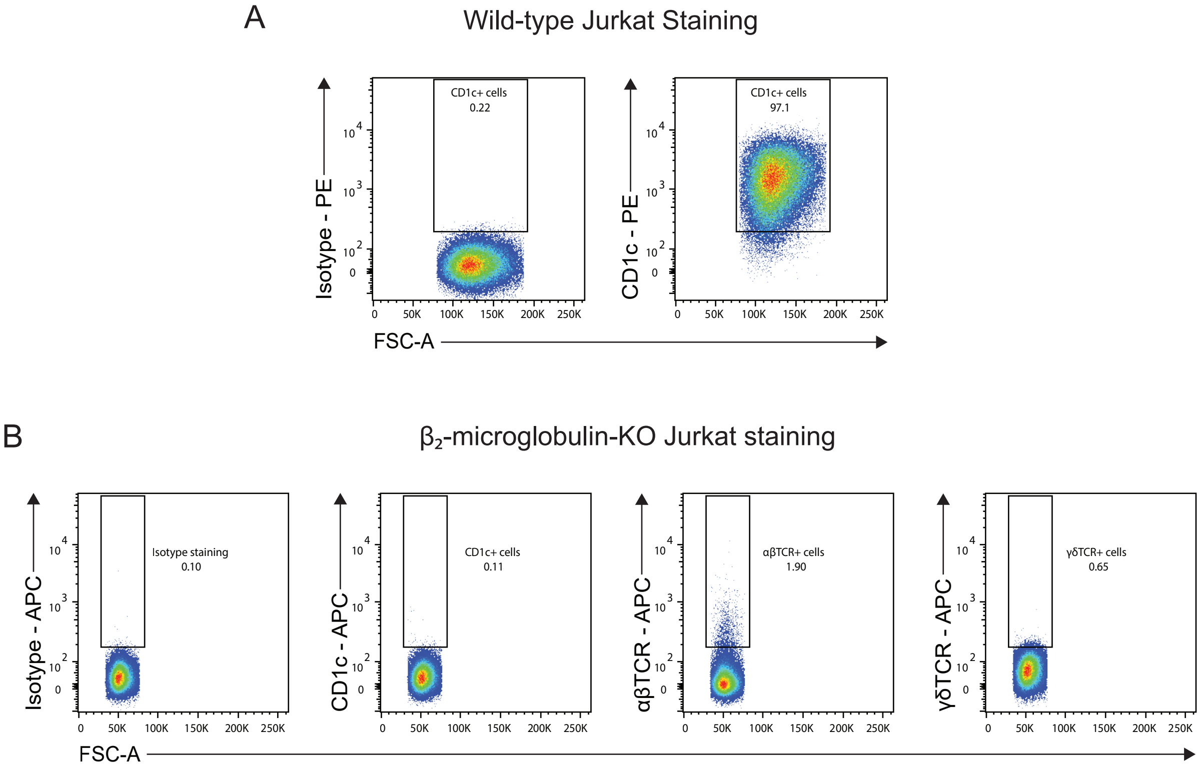


**Fig.S4.** (A) Flow cytometry dot plots showing high expression of CD1c on wild-type JRT3.5 Jurkat T-cells. (B) Flow cytometry dot plots showing the absence of CD1c and TCR expression on β2-microglobulin knock-out JRT3.5 Jurkat T-cells.

### Figure S5

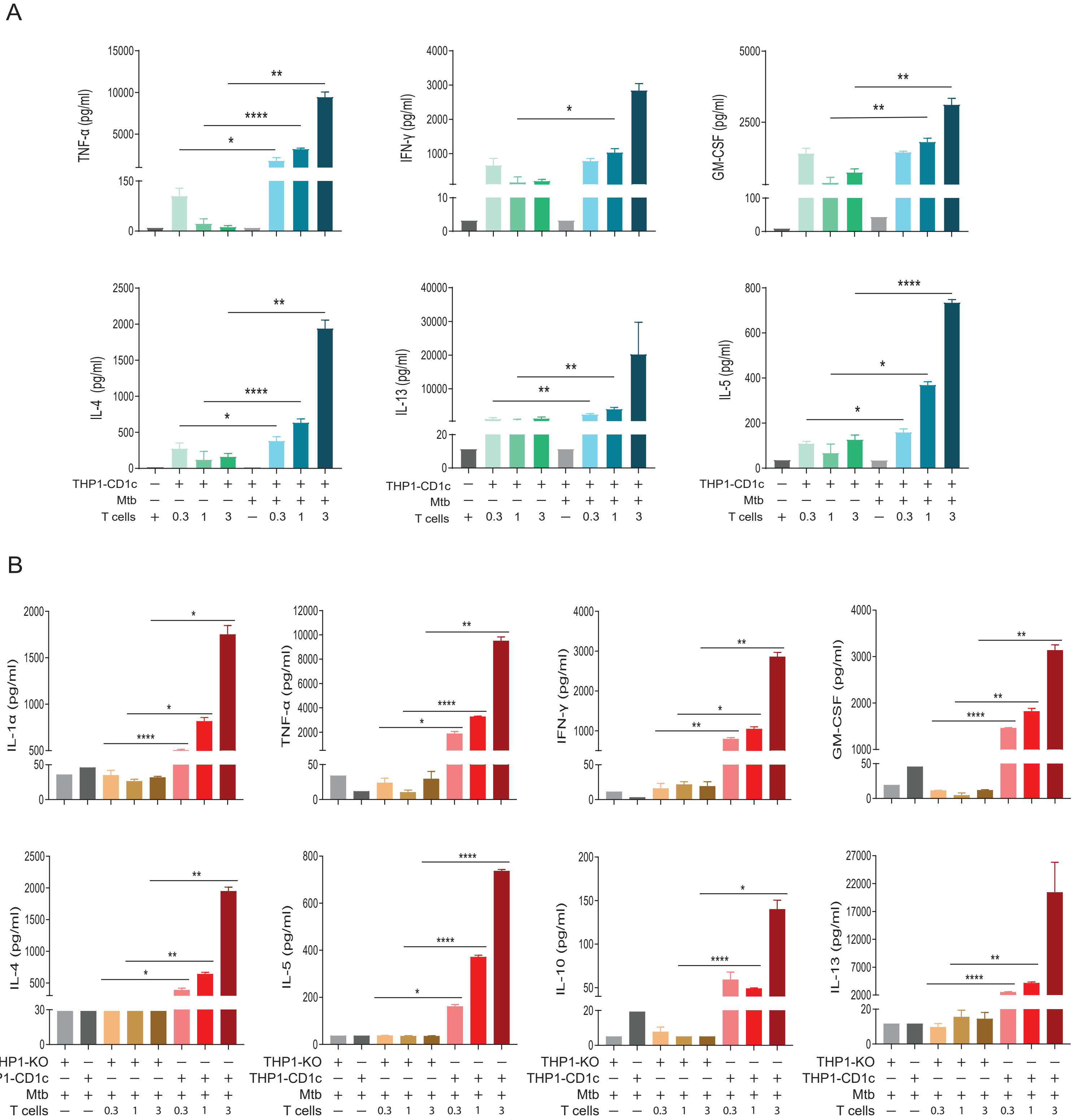


**Fig.S5.** (**A**) CD1c-autoreactive T-cells secrete cytokines in response to uninfected THP1-CD1c APCs in an autoreactive manner. Importantly, CD1c-autoreactive T-cells release significantly higher concentrations of cytokine when cultured with Mtb-infected THP1-CD1c APCs. (**B**) CD1c-autoreactive T-cells secrete diverse cytokines in a CD1c dependent manner. CD1c-autoreactive T-cells release cytokines when cultured with Mtb-infected THP1-CD1c APCs, but not when were cultured with Mtb-infected THP1-KO APCs. Data are representative of two independent experiments, each preformed in triplicate. * P < 0.05; ** P < 0.01, **** P < 0.0001 (Two-way ANOVA).

### Figure S6

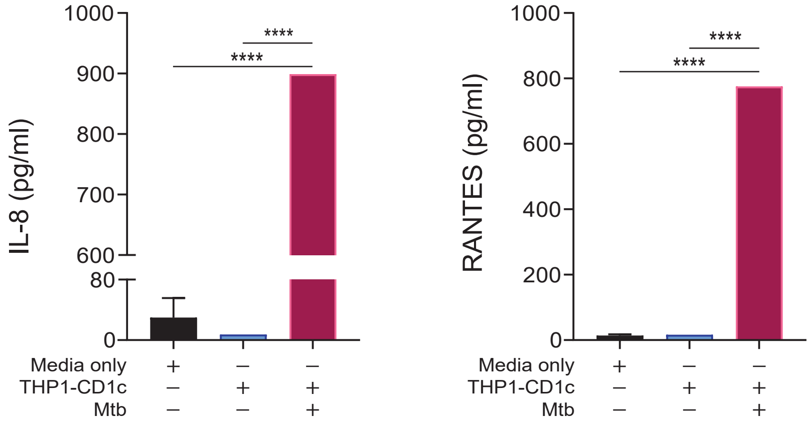


**Fig.S6.** Mtb-infected THP1-CD1c APCs secrete chemokines IL-8 and RANTES (pink bars), in comparison to media only (black bars), and uninfected THP1-CD1c APCs (blue bars). Data are representative of two independent experiments, each preformed in triplicate. **** P < 0.0001 (Two-way ANOVA).

### Figure S7

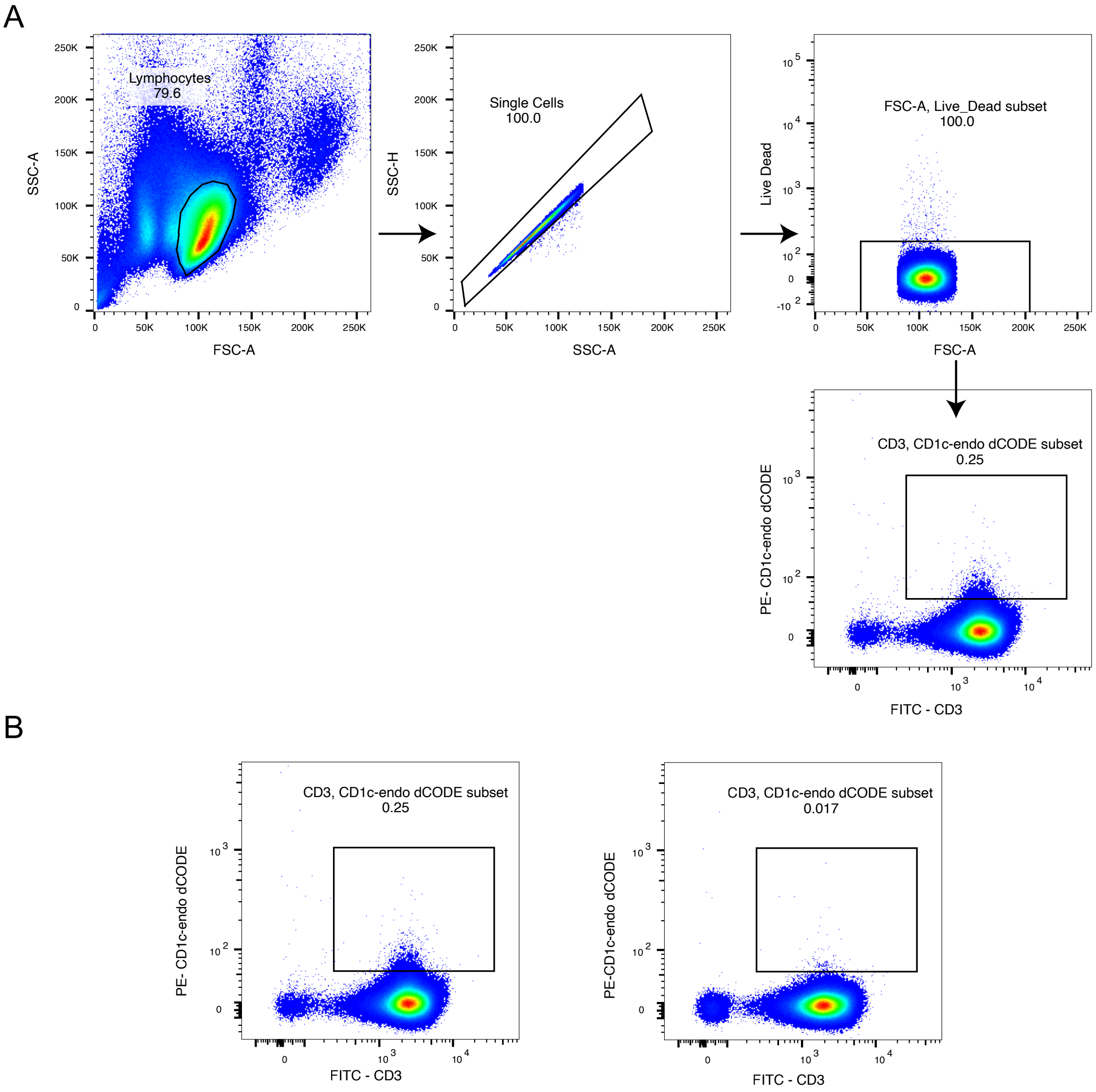


**Fig.S7.** (A) Representative flow cytometry gating strategy used to isolate CD3+ T-cells stained with CD1c-endo dCODE dextramers. (B) Representative plots from two donors showing distinct CD3 +CD1c endo positive and CD3+CD1c-endo negative T-cell populations. Percentages indicate the proportion of positive cells within the CD3+ gate.

### Figure S8

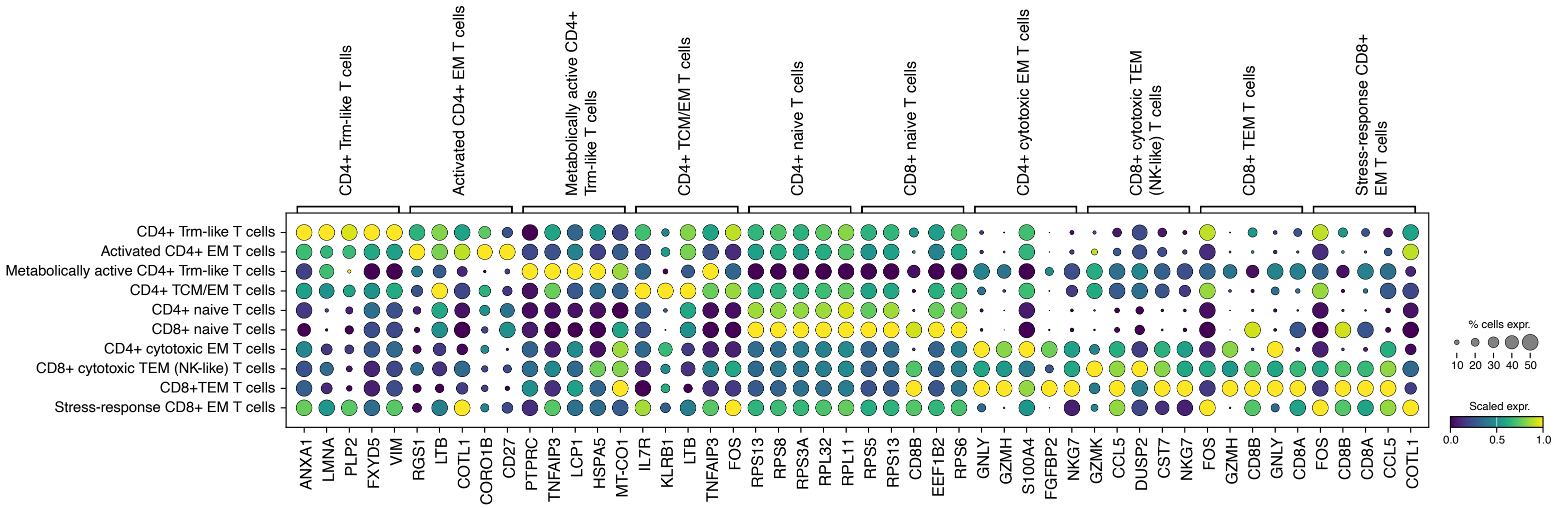

**Fig.S8.** Dot plot showing the top 5 ranked marker genes for each annotated T-cell population, derived from all high-quality T-cells (n = 11,804). Dot size represents the proportion of cells expressing each gene within the corresponding cluster, while colour indicates the scaled average expression (z-score). This visualization highlights the transcriptional signatures that define distinct T-cell subsets, including naïve, memory, effector, and metabolically active populations.

### Figure S9

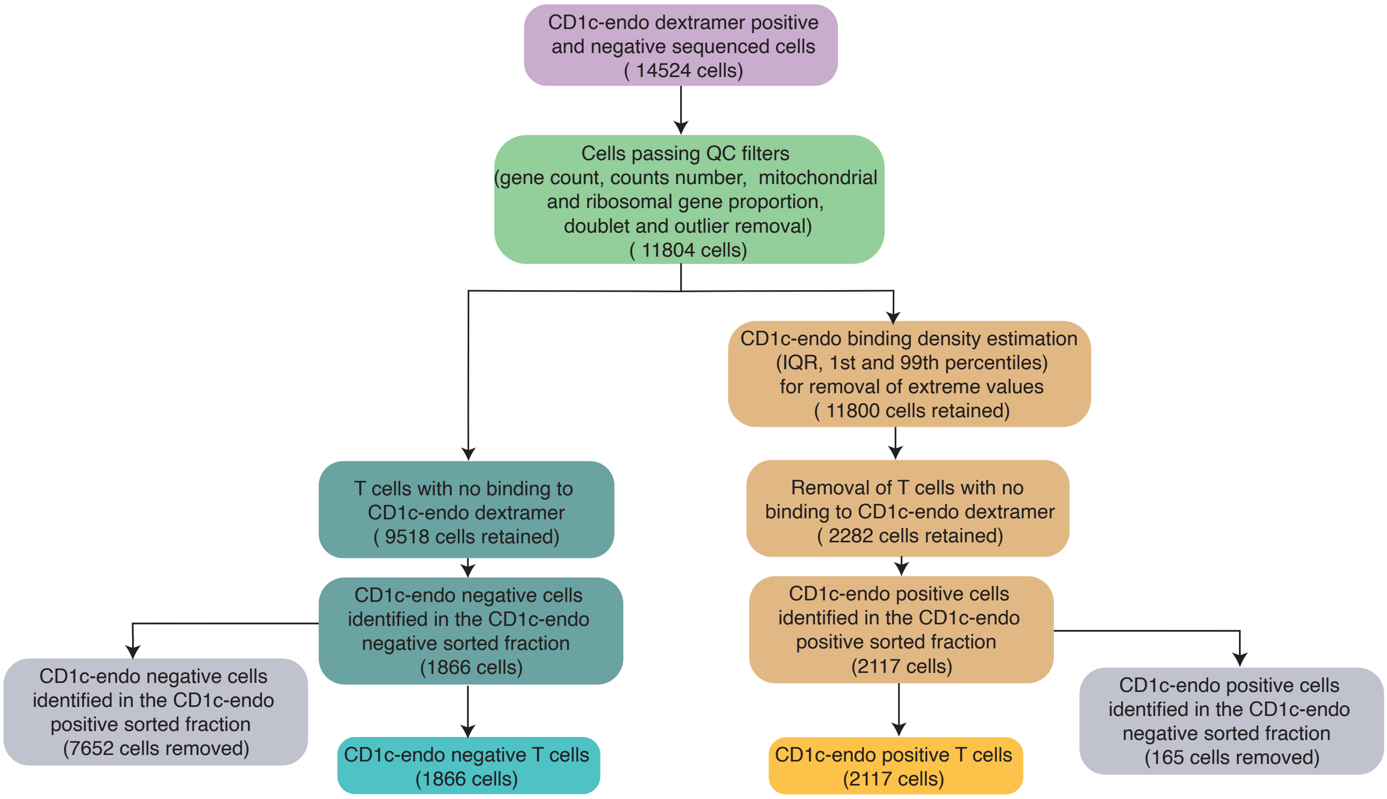


**Fig.S9**. Filtering strategy for identifying CD1c-endo dextramer-bound T-cells in scRNA-seq data from sorted populations: A total of 14,524 cells were initially profiled by scRNA-seq from dextramer-sorted fractions. Following standard quality control, including thresholds for gene and UMI counts, mitochondrial and ribosomal content, doublet detection, and outlier removal, 11,804 high-quality cells were retained. CD1c-endo-negative cells (n = 9,518) were defined by the absence of detectable dextramer signal and further stratified by their sort gate: 7,652 originated from the CD1c-endo-positive gate and 1,866 from the CD1c-endonegative gate. To minimise false negatives, only the 1,866 cells from the negative sort gate were retained as confidently CD1c-endo-negative. To define CD1c-endopositive cells, dextramer signal intensities were modelled using kernel density estimation (KDE) and quantile-based filtering (1st and 99th percentiles, IQR). Cells with signal below the lower bound (Q1 – 1.5×IQR) or extreme values were excluded, yielding 11,800 cells. Of these, 2,288 cells with no binding signal were removed. Among the remaining 2,282 CD1c-endo-positive candidates, 2,117 were from the CD1c-endo-positive sort gate and 165 from the negative gate. To ensure specificity, only the 2,117 cells from the positive sort gate were retained as confidently CD1c-endo-positive. This combined quality control and gating strategy enabled robust identification of CD1c-endo-binding T-cells in the single-cell dataset.

**Table S1. Primer sequences used for targeted single-cell TCR sequencing.**

Sequences are shown for the oligo-dT primer, universal forward and reverse primers, targeted forward primer containing the 8-bp well barcode, and reverse primers specific for TRAC, TRBC, TRGC and TRDC constant regions.

| | |
|---|---|
| Oligo-dT | GGAATTCGACCGAGTACACATAAGGT30VN |
| Universal Forward | ACGTCACTCACTCGTAGCCTAAGCAGTGGTATCAACGCAGAGT |
| Universal Reverse | GGAATTCGACCGAGTACACATAAGG |
| Targeted Forward | 8 bp well barcode-ACGTCACTCACTCGTAGCCT |
| Targeted Reverse TRAC | GAGTCTCTCAGCTGGTACACGG |
| Targeted Reverse TRBC | AGTGTGGCCTTTTGGGTGTG |
| Targeted Reverse TRGC | GGTATGTTCCAGCCTTCTGGAG |
| Targeted Reverse TRDC | GGGAGAGATGACAATAGCAGGATC |
